# Dramatically diverse *S. pombe wtf* meiotic drivers all display high gamete-killing efficiency

**DOI:** 10.1101/728535

**Authors:** María Angélica Bravo Núñez, Ibrahim M. Sabbarini, Michael T. Eickbush, Yue Liang, Jeffrey J. Lange, Aubrey M. Kent, Sarah E. Zanders

**Author notes:** Correspondence to: Sarah E. Zanders, Stowers Institute for Medical Research, 1000 E 50^th^ Street, Kansas City, MO 64110.

## Abstract

Meiotic drivers are selfish genetic loci that force their transmission into more than 50% of the viable gametes made by heterozygotes. Meiotic drivers are known to cause infertility in a diverse range of eukaryotes and are predicted to affect the evolution of genome structure and meiosis. The *wtf* gene family of *Schizosaccharomyces pombe* includes both meiotic drivers and drive suppressors and thus offers a tractable model organism to study drive systems. Currently, only a handful of *wtf* genes have been functionally characterized and those genes only partially reflect the diversity of the *wtf* gene family. In this work, we functionally test 22 additional *wtf* genes. We identify eight new drivers that share between 30-90% amino acid identity with previously characterized drivers. Despite the vast divergence between these genes, they generally drive into >85% gametes when heterozygous. We also find three new *wtf* genes that suppress drive, including two that also act as autonomous drivers. Additionally, we find that *wtf* genes do not underlie a weak (64%) transmission bias caused by a locus or loci on chromosome 1. Finally, we find that some Wtf proteins have expression or localization patterns that are distinct from the poison and antidote proteins encoded by drivers and suppressors, suggesting some *wtf* genes may have non-meiotic drive functions. Overall, this work expands our understanding of the *wtf* gene family and the burden selfish driver genes impose on *S. pombe*.

**Article Summary:** During gametogenesis, the two gene copies at a given locus, known as alleles, are each transmitted to 50% of the gametes (e.g. sperm). However, some alleles cheat so that they are found in more than the expected 50% of gametes, often at the expense of fertility. This selfish behavior is known as meiotic drive. Some members of the *wtf* gene family in the fission yeast, *Schizosaccharomyces pombe*, kill the gametes (spores) that do not inherit them, resulting in meiotic drive favoring the *wtf* allele. Other *wtf* genes act as suppressors of drive. However, the *wtf* gene family is diverse and only a small subset of the genes has been characterized. Here we analyze the functions of other members of this gene family and found eight new drivers as well as three new suppressors of drive. Surprisingly, we find that drive is relatively insensitive to changes in *wtf* gene sequence as highly diverged *wtf* genes execute gamete killing with similar efficiency. Finally, we also find that the expression and localization of some Wtf proteins are distinct from those of known drivers and suppressors, suggesting that these proteins may have non-meiotic drive functions.

## Introduction

During meiosis, diploid cells divide to produce haploid gametes (e.g. sperm). This process is generally fair in that each parental allele of a gene is represented at an equal ratio in the gametes (1). However, many eukaryotic genomes contain ‘selfish’ elements that bias their own transmission into the viable gametes generated by a heterozygote (2–4). These loci are known as meiotic drivers and they can directly and indirectly reduce fitness through a number of mechanisms (reviewed in (5, 6)).

Meiotic drivers are in conflict with unlinked genes that do not gain a transmission advantage but must bear the fitness burdens imposed by these selfish elements (7–10). This genetic conflict is thought to favor the emergence of unlinked allele variants that can suppress the effects of drivers. In turn, variants of drivers that can evade this suppression will have a selective advantage (4, 8, 11). The conflict between drivers and suppressors is predicted to lead to the rapid evolution of both sets of genes. This is analogous to the genetic arms race observed between viruses and host immune systems (12, 13). Arms races caused by meiotic drivers are predicted to affect the evolution of gametogenesis and genome structure (14, 15).

Empirical analyses of meiotic drive/suppressor systems have traditionally been limited by the complexity of most known drive systems. However, the *wtf* (***<u>w</u>****ith **<u>tf</u>***) gene family of the fission yeast *Schizosaccharomyces pombe* offers an opportunity to study meiotic drive in a highly tractable model system. Some *wtf* genes are meiotic drivers and others act as suppressors of drive (16–18). Characterized *wtf* drivers kill the meiotic products (spores) that do not inherit them from a *wtf+/wtf-* heterozygous diploid by producing two proteins described as the Wtf^poison^ and Wtf^antidote^. These proteins are encoded on largely overlapping transcripts, but the antidote message includes a 5’ exon (exon 1) not found in the poison message (Figure 1A). The poison protein localizes throughout the spore sac (ascus) and acts on every spore in an indiscriminate fashion. The antidote protein, however, is highly enriched within the *wtf*+ spores, rescuing them from destruction by the poison (16, 17). Similarly, the known drive-suppressing *wtf* genes produce a protein that mimics the antidote of a driver, rescuing the spores that inherit the suppressor from the driver’s poison (18).

**Figure 1.**
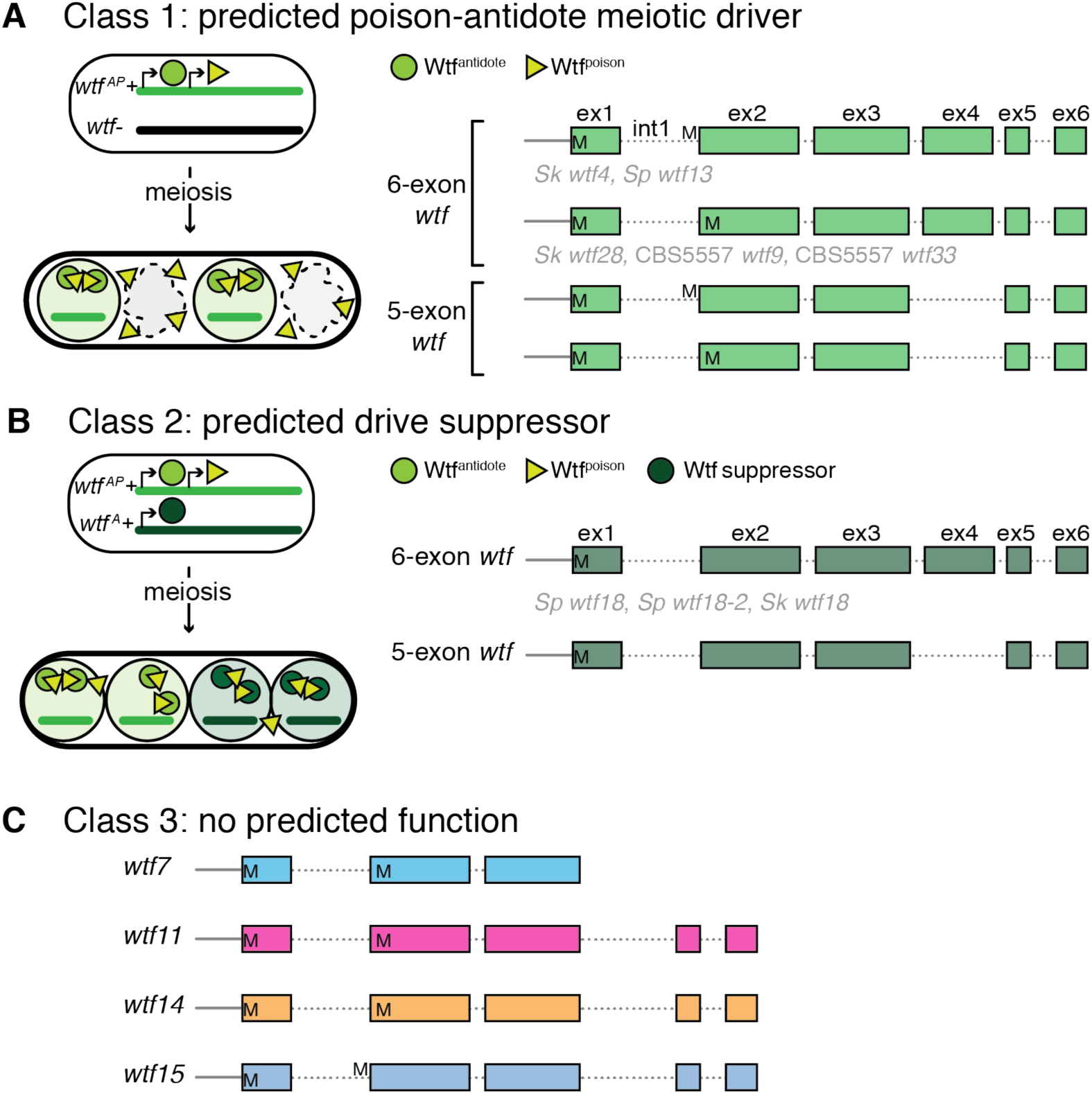
Classification of the *wtf* gene family. (A) Model of how known *wtf* meiotic drivers kill the spores that do not inherit them from a *wtf+/wtf-* diploid. *wtf* meiotic drivers make a trans-acting poison and an antidote that is enriched in the spores that inherit the *wtf* allele. Class 1 contains both 6- and 5-exon *wtf* genes. The previously characterized *wtf* meiotic drivers are shown in gray underneath the gene structure (16–18). (B) Model for how antidote-only *wtf* genes can work as suppressors of *wtf* meiotic drivers. The spores that inherit the *wtf* antidote-only gene are protected from a specific Wtf^poison^. Class 2 contains the predicted antidote-only *wtf* genes. The previously characterized 6-exon *wtf* suppressors are listed underneath the gene structure (18). (C) Gene structure of class 3 genes. These genes have no predicted function. M’s (methionine) represent potential translational start sites.

The *wtf* gene family contains 25 members in the *S. pombe* reference strain, *L972* (referred to here as ‘*Sp*’) (19, 20). This gene family is rapidly evolving and other *S. pombe* isolates, in which the *wtf* genes have been assembled, contain up to 38 members (17, 21). Currently, only a handful of these genes have been characterized (16–18). Although the characterized genes have provided mechanistic insight, they do not represent the extensive diversity within the *wtf* gene family. It is currently unclear if the sequence diversity of *wtf* genes corresponds with phenotypic diversity or if some *wtf* genes could have a function outside of drive.

To aid in the functional characterization of the *wtf* gene family, we previously sorted them into three classes. **Class 1** genes are predicted to be meiotic drivers (Figure 1A). These genes contain two potential transcriptional and translational start sites that could be used to encode distinct poison and antidote proteins. **Class 2** genes are predicted drive suppressors that appear to encode only Wtf^antidote^ proteins (Figure 1B). **Class 3** includes four diverse genes (*wtf7, wtf11, wtf14,* and *wtf15)* that are grouped together only because they are unlike all other *wtf* genes (Figure 1C). These genes also have two potential translational start sites, but unlike class 1 genes, they produce only one major transcript type (22). Additionally, class 3 *wtf* genes are less diverged within isolates of *S. pombe* than the *wtf* genes in class 1 and class 2 (21). These *wtf* genes have no predicted function.

Here we use a combination of genetics and cell biology approaches to functionally analyze 22 previously untested *wtf* genes. We find that a tremendous breadth of Wtf protein sequences can build highly efficient poison-antidote drivers and drive suppressors. Moreover, we discover that some *wtf* genes can simultaneously act as drivers and as suppressors of other drive loci. Surprisingly, we also find that *wtf* genes do not underlie a previously identified but unmapped locus on chromosome 1 that causes an allele transmission bias in the viable spores. Finally, we discover that the potential cellular roles of *wtf* family members extend beyond gametogenesis.

## Results

### Class 1 contains autonomous meiotic drivers

To test the ability of class 1 and class 2 *wtf* genes to drive autonomously, we cloned untested genes representing unique subclasses from four *S. pombe* isolates: *Sp*, *S. kambucha* (*Sk*), FY29033, and CBS5557 (Supplemental Figure 1). We then assayed the ability of these *wtf* genes to drive in one or more ectopic strain backgrounds (e.g. testing *Sk wtf* genes in *Sp*). We used ectopic strain backgrounds because drive occurs in heterozygotes and it is simpler to insert genes into *S. pombe* than it is to delete the repetitive *wtf* genes at their endogenous loci. Additionally, drivers may be less likely to be suppressed in an ectopic background (23).

To do these analyses, we first linked each *wtf* gene under the control of its endogenous promoter to a selectable drug resistance marker (*kanMX4* or *hphMX6*) and integrated them into the *ade6* locus (of either *Sp* and/or *Sk).* We then crossed each haploid carrying a *wtf* gene of interest to a wild-type strain from the same background to generate hemizygous (*wtf::*drug^R^/*ade6*+) diploid strains. We induced the hemizygous diploids to undergo meiosis and then assayed the presence of each *wtf* gene of interest in the viable spore population using the linked selectable markers. Genes capable of autonomous drive are expected to be overrepresented in the viable progeny (>50%).

Most genes within class 1 have six exons, including all previously described *wtf* drivers. Class 1 also includes two 5-exon *wtf* genes (Figure 1A, Supplemental Figure 1). We tested ten 6-exon and two 5-exon *wtf* genes from class 1 (Figure 2A, diploids 1-16). We found that seven of the 6-exon genes and one 5-exon gene exhibited significant drive in at least one strain background (Figure 2A). These genes are: *Sk wtf9, Sk wtf19, Sk wtf30, Sk wtf33, Sp wtf19,* FY29033 *wtf36,* FY29033 *wtf18,* and FY29033 *wtf35*. Interestingly, these genes are incredibly diverse and share as little as 30% pairwise amino acid identity (Supplemental Figure 2A). Moreover, all of the drivers except one (*Sp wtf19*) drive into >90% of the progeny when tested under the same conditions (16, 18). These results demonstrate that a remarkably wide range of proteins can execute spore-killing similarly well. Interestingly, the 5-exon FY29033 *wtf35* meiotic driver lacks sequences homologous to what is exon 4 in the other known 6-exon drivers. In addition, the gene lacks the 7-amino acid repeat that is found in between 1-4 copies in the last exon of all other drivers (21). The fact that FY29033 *wtf35* can still drive suggests these sequences can be dispensable for drive.

**Figure 2.**
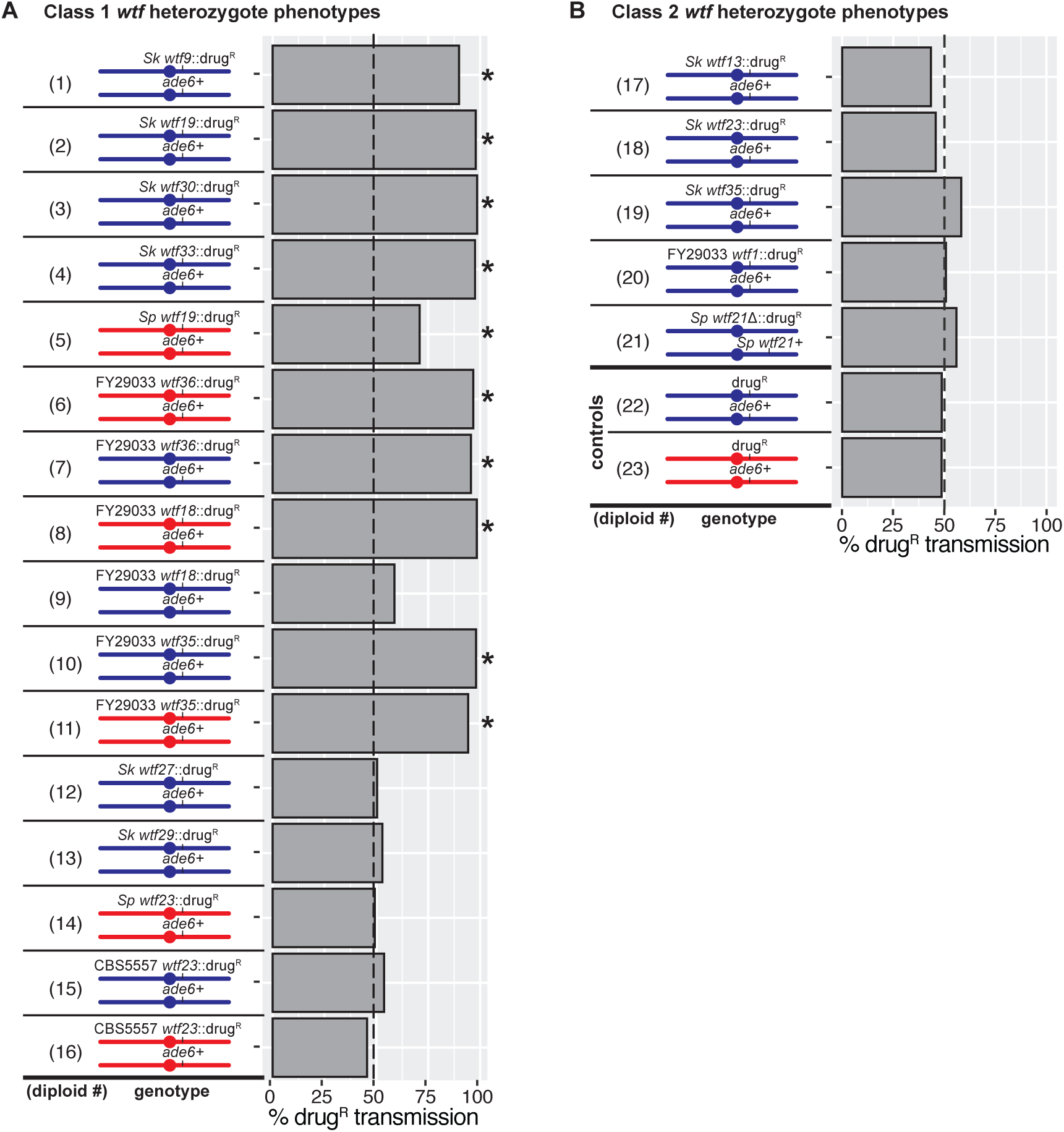
Phenotypes of *wtf+/wtf-* diploids. Allele transmission of (A) class 1 *wtf* gene and (B) class 2 *wtf* gene heterozygotes. We used the *kanMX4* or *hphMX6* drug resistance markers (drug^R^) linked to the *wtf* allele of interest and the *ade6* marker to follow allele transmission. For diploids 1-20, we excluded the spores that had inherited both *ade6+* and drug^R^ markers as these are likely aneuploid or diploid. For diploid 21, allele transmission was assayed using only the drug marker. Diploids 1-4,7,9,10,12,13,15, and 17-20 were compared to control diploid 22; diploids 5,6,8,11,14, and 16 were compared to the control diploid 23. *Sp* chromosomes are depicted in blue and *Sk* chromosomes are shown in red. * indicates a p-value<0.02 (G-test). We genotyped at least 200 haploid offspring for each diploid. We tested some of the *wtf* transgenes using multiple independent strains (i.e. one in which the transgene was marked with *kanMX4* and one marked with *hphMX6*) and we present the combined data. The complete raw data are presented in Supplemental Table 1.

The four class 1 genes that did not exhibit drive in our tests were *Sk wtf27, Sk wtf29, Sp wtf23,* and CBS5557 *wtf23* (Figure 2A, diploids 12-16). CBS5557 *wtf23* completely lacks a specific 11-amino acid repeat within exon three that can be found in 2-7 copies in all characterized drivers except FY29033 *wtf35*, which contains only four amino acids of the repeat. We found no sequence features in *Sk wtf27, Sk wtf29,* or *Sp wtf23* that distinguish these genes from the confirmed drivers. While it is possible our results reflect that these four genes are not capable of driving, it is also possible that these genes are suppressed in the background in which we tested them. We previously saw background-specific suppression of the *Sp wtf13* driver, and similar suppression explains why we observed drive of FY29033 *wtf18* in the *Sk*, but not in the *Sp* background (see below) (18). Overall, our analyses are consistent with class 1 containing a wide diversity of autonomous meiotic drive genes.

The class 2 *wtf* genes are either known or predicted antidote-only drive suppressors (Figure 1B). Like the class 1 genes, class 2 contains both 5-exon and 6-exon genes, although only 6-exon genes have been previously tested (16–18). Interestingly, class 2 also contains *Sp wtf21*, which was previously reported to be an essential gene because a heterozygous mutant (*Sp wtf21/wtf21Δ*) transmitted only the wild-type *Sp wtf21* allele to viable gametes (24).

We first tested four class 2 genes (*Sk wtf13, Sk wtf23*, *Sk wtf35,* and FY29033 *wtf1*) using the same approach described above for class 1 genes. As expected for genes predicted to produce only antidotes, we found that none of the genes tested exhibited drive in an ectopic strain background (Figure 2B, diploids 17-20). We also revisited the idea that *Sp wtf21* is an essential gene (24). We found that we could generate a deletion of *Sp wtf21* in a haploid strain, indicating the gene is not essential in that strain background. Moreover, we generated an *Sp wtf21/wtf21Δ* heterozygote and did not observe drive (Figure 2B, diploid 21).

Finally, we tested if we could observe autonomous meiotic drive by the class 3 *wtf* genes (*wtf7, wtf11, wtf14,* and *wtf15*) in *Sp*. Unlike the other *wtf* genes (class 1 and class 2), each natural isolate has a clear ortholog of each class 3 *wtf* gene (21). We therefore had to use a different strategy to test if these genes could drive. Instead of introducing the genes into an ectopic strain background, we assayed whether these genes could drive when heterozygous at their endogenous loci. We deleted *wtf7* and *wtf11* individually, and *wtf14* and *wtf15* together as they are adjacent to each other. In diploids heterozygous for any of these *wtf* gene deletions, the wild-type alleles were transmitted to ∼50% of the viable spores (Supplemental Figure 3A). This indicates that these genes cannot drive or that they are suppressed in the *Sp* background. We also assayed a homozygous deletion strain lacking all of the class 3 genes. We observed no fertility defects, indicating these genes are not required for sexual reproduction (Supplemental Figure 3B). We also observed no growth defects in haploids lacking the class 3 genes (Supplemental Figure 3C).

### *wtf* genes in both class 1 and class 2 can act as suppressors of drive

Previous work identified alleles of *wtf18* in *Sp* and *Sk* as suppressors of meiotic drive, but we wanted to test if other *wtf* genes could also act as drive suppressors (18). Within drivers and their known suppressors, the Wtf^antidote^ proteins share high levels of amino acid identity with the poisons they neutralize. This similarity appears to be particularly important within the C-terminus (18). We used this knowledge to guide our search for other drive suppressors. We found that the residues encoded in the last three exons of the class 2 gene FY29033 *wtf1* share >99% identity to those in the FY29033 *wtf35* driver (Figure 3A). We therefore predicted that FY29033 *wtf1* could be a suppressor of the FY29033 *wtf35* driver. To test this, we made diploids heterozygous for both alleles (FY29033 *wtf35/*FY29033 *wtf1*) and assayed transmission of each allele. FY29033 *wtf35* was no longer able to drive in the presence of FY29033 *wtf1* in the *Sp* background (Figure 3C, compare diploid 10 to 24). Additionally, FY29033 *wtf1* was able to rescue the fertility defect caused by FY29033 *wtf35* (Figure 3C). These results demonstrate that FY29033 *wtf1* is a suppressor of FY29033 *wtf35*.

**Figure 3.**
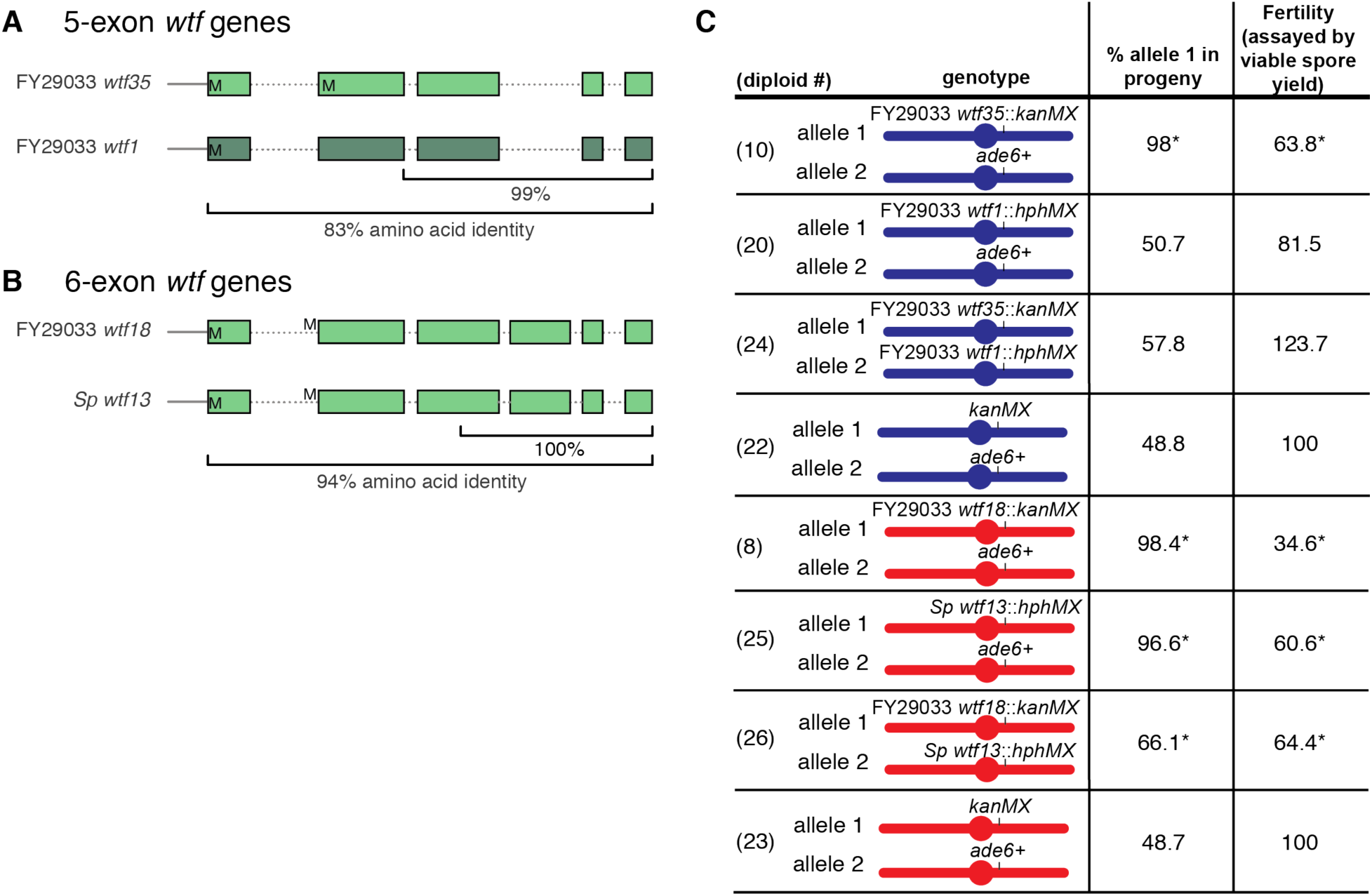
*wtf* genes can be both drivers and suppressors of drive. Cartoons illustrating the similarity between (A) FY29033 *wtf35* and FY29033 *wtf1* and (B) FY29033 *wtf18* and *Sp wtf13*. M’s (methionine) represent the translational start sites. (C) Allele transmission and fertility phenotypes of *Sp* (blue) and *Sk* (red) diploids with the indicated transgenes integrated at *ade6.* Allele transmission was determined by following the genetic markers (*ade6* and drug resistance) linked to each allele. Diploids 10, 20, and 24 were compared to control diploid 22, while diploids 8, 25, and 26 were compared to control diploid 23. The data for diploid 25 was previously published in (18). We normalized the fertility values to the control diploid and reported them as a percentage. * indicates p-value of <0.05 (G-test [allele transmission] and Wilcoxon test [fertility]). We genotyped more than 200 haploid spores for each diploid. Spores that inherited both markers at *ade6* (Ade+ G418^R^, Ade+ HYG^R^, or G418^R^ HYG^R^) were excluded from the analyses as they are likely aneuploid or diploid. We present the complete raw data in Supplemental Tables 1 and 2.

To broaden our search for suppressors, we examined why FY29033 *wtf18* drives in *Sk,* but not in *Sp.* Specifically, we looked for *wtf* genes in *Sp* that could work as a suppressor of FY29033 *wtf18.* We noticed that the C-terminal region (216 amino acids) of FY29033 *wtf18* is identical to that of the *Sp wtf13* meiotic driver (Figure 3B). This suggested that the *Sp* Wtf13^antidote^ could potentially neutralize the FY29033 Wtf18^poison^ and *vice versa*. To test this, we generated FY29033 *wtf18*::*kanMX4/Sp wtf13*::*hphMX6* heterozygotes in an *Sk* strain background and assayed their phenotypes. We observed that drive of both genes was suppressed in the heterozygote relative to hemizygotes containing only one of the drivers (Figure 3, compare diploids 8 and 25 to diploid 26).

We still observed drive of the FY29033 *wtf18* allele in the FY29033 *wtf18/Sp wtf13* heterozygous diploid, suggesting the Wtf13^antidote^ is only partially effective against the Wtf18^poison^. Additionally, the level of drive of FY29033 *wtf18* we observed is sufficient to explain the decrease in fertility of the diploid (64.4% of wild-type, Figure 3C, diploid 26). These results show that *wtf* drivers can also function as suppressors of each other.

### *wtf* genes are not responsible for a meiotic allele transmission bias favoring *Sk* chromosome 1

We previously detected a weak allele transmission bias that favored *Sk* chromosome 1 in the viable progeny of *Sp/Sk* hybrid diploids (25). We reexamined this observation here with hybrid diploids that are heterozygous for *Sp* and *Sk* copies of chromosome 1, but are homozygous for *Sk* chromosomes 2 and 3. These hybrids were also unable to initiate meiotic recombination due to deletion of *rec12* (*SPO11* homolog), which is required for programmed meiotic double-stranded break formation. The *rec12* deletion ensured chromosome-wide linkage on chromosome 1, allowing us to monitor transmission of the unmapped locus into the spore progeny (Supplemental Figure 4). Consistent with our previous observations, we observed 64.3% of the viable spores generated by the hybrids inherited *Sk* chromosome 1. There is only one *wtf* locus, *wtf1,* on both *Sp* and *Sk* chromosome 1. We found that the transmission of *Sk* chromosome 1 was not significantly different after deleting *Sk wtf1* (Supplemental Figure 4B). This result is consistent with the presence of a non-*wtf* driver on *Sk* chromosome 1. However, it is also possible that Dobzhansky-Muller incompatibilities between *Sp* chromosome 1 and *Sk* chromosomes 2 and/or 3 explain the transmission bias (26).

### High amino acid identity is crucial for Wtf poison and antidote specificity

We next wanted to use our expanded knowledge of drivers and suppressors to further refine our understanding of how similar antidote proteins must be to the poisons they neutralize. All known suppressors have >97% identity in the amino acids encoded in the last two exons to the drivers they suppress, but it is unclear if less similar Wtf proteins could also work. We reasoned that genes that drive in *Sp* must not be suppressed by any endogenous *Sp* genes. We therefore compared the similarity of genes that drive in *Sp* to every *Sp wtf* gene that failed to suppress them (i.e. transcribed *wtf* genes) (22). We found that the endogenous *Sp* Wtf proteins and the ectopic drivers share between 43-86% pairwise identity within the residues encoded in the final two exons (43-69 amino acids) (Supplemental Figure 5A and 5B). This analysis is consistent with the notion that high amino acid identity within the C-terminal region of the Wtf antidote and poison proteins (87-100%) is important for specificity (Supplemental Figure 5A and 5C).

**Figure 5.**
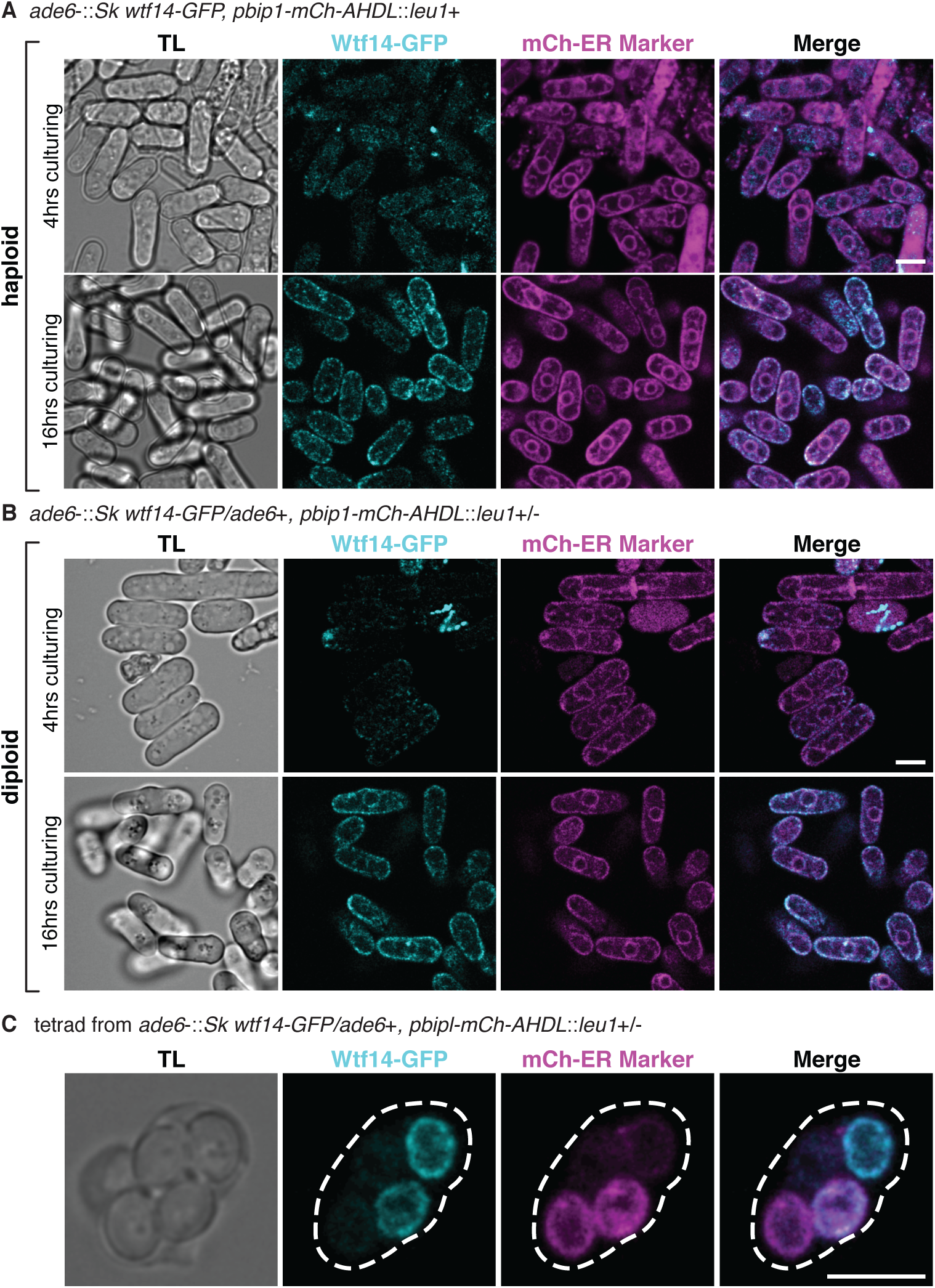
*Sk wtf14-GFP* is broadly expressed and localizes to the perinuclear and cortical endoplasmic reticulum (ER). Representative images of *Sk* Wtf14-GFP (cyan) and ER marker mCherry-AHDL (magenta) in (A) haploids and (B) diploids after 4 hours and 16 hours of culturing are shown. The imaged diploids were heterozygous for both *Sk wtf14-GFP* and *pbip1-mCh-AHDL*. (C) A tetrad generated by a diploid heterozygous for *Sk wtf14-GFP* and *pbip1-mCh-AHDL* is shown. We linearly unmixed the images in A and B (Materials and Methods, Supplemental Figure 10). We adjusted the brightness and contrast differently for each image and smoothed them using Gaussian blur. The scale bar represents five microns. TL, transmitted light.

### Class 3 *wtf* genes are not required for drive of *Sk wtf4*

We previously analyzed the sequences of the class 3 *wtf* genes across four isolates of *S. pombe* (*Sp*, *Sk,* FY29033, and CBS5557) and found that *wtf7*, *wtf11,* and *wtf15* show signatures of positive selection (21). This suggests that selection is favoring novelty and led us to speculate that these genes could be involved in meiotic drive. Given that we failed to observe drive of these class 3 genes (Supplemental Figure 3A), we thought they could perhaps be modifying drive of other *wtf* genes. All class 3 *wtf* genes are linked to drive loci so we predicted that they could facilitate drive, as they would also gain a transmission advantage from a linked driver (8).

We tested this idea by generating diploids hemizygous for the *Sk wtf4* meiotic driver (*Sk wtf4*::*kanMX4*/*ade6+*) and lacking all four of the class 3 *wtf* genes. *Sk wtf4* was found in nearly 100% of the viable spores generated by this quadruple deletion mutant (Supplemental Figure 3D, compare diploid 30 to 31). These results show that *wtf7*, *wtf11*, *wtf14*, and *wtf15* are not required to facilitate drive of *wtf* meiotic drivers.

### Localization of class 3 *Sk* Wtf proteins in *Sp*

We next decided to take a more agnostic approach to look for possible functions of Wtf7, Wtf11, Wtf14, and Wtf15 by investigating their expression and localization in cells. We analyzed a published data set from a proteomics meiotic time course study (27) and found that peptides from Wtf11, Wtf14, and Wtf15, but not Wtf7, were all detected during meiosis in three replicate experiments (Supplemental Figure 6). Wtf11 and Wtf15 were detected in one or more timepoints taken after the first meiotic division. Interestingly, the levels of Wtf14 remained steady throughout meiosis (Supplemental Figure 6). These patterns are both unlike those of class 1 and class 2 Wtf proteins which increase in abundance as meiosis progresses (18, 27).

We also tagged each of the class 3 genes under the control of their endogenous promoters (cloned from the *Sk* isolate) with GFP at the C-terminus and integrated them at the *ade6* locus of *Sp*. We then imaged haploid and diploid cells during logarithmic cell growth and stationary phase. We also imaged cells undergoing meiosis and tetrads. In contrast to the proteomics data, we were not able to detect any GFP fluorescence under any conditions in cells containing the *wtf11-GFP* allele (Supplemental Figure 7) (27). The reasons for this are unclear, but it could be due to our tag generating a null allele. We also did not detect Wtf7-GFP or Wtf15-GFP in vegetatively growing haploids or diploids (Supplemental Figure 8). However, we did observe GFP fluorescence in tetrads produced by diploids heterozygous for *wtf7-GFP* and tetrads produced by *wtf15-GFP* heterozygotes (Figure 4 and Supplemental Figure 9). This late expression of Wtf15 is consistent with the proteomics data set (Wtf7 was not detected by proteomics) (27). In tetrads generated by diploids with one tagged copy of Wtf7 or Wtf15, we observed that the signal was greatly enriched in two of the four spores (Figure 4). We speculate that the two spores with bright GFP signal are those that inherited the tagged allele. This would suggest that *wtf7-GFP* and *wtf15-GFP* are both expressed after spore individualization, similar to the antidotes of *wtf* drivers and suppressors (16, 18). Wtf15-GFP exhibited a diffuse localization pattern that largely filled the spores (Figure 4B and Supplemental Figure 9B). The localization pattern of Wtf7-GFP varied in different spores. In some instances, Wtf7-GFP made a ring-like structure next to Nsp1-mCherry (nucleoporin marker) (Figure 4A and Supplemental Figure 9A). However, we also observed other cases where Wtf7-GFP seemed to be clustered within the spore (Figure 4A).

**Figure 4.**
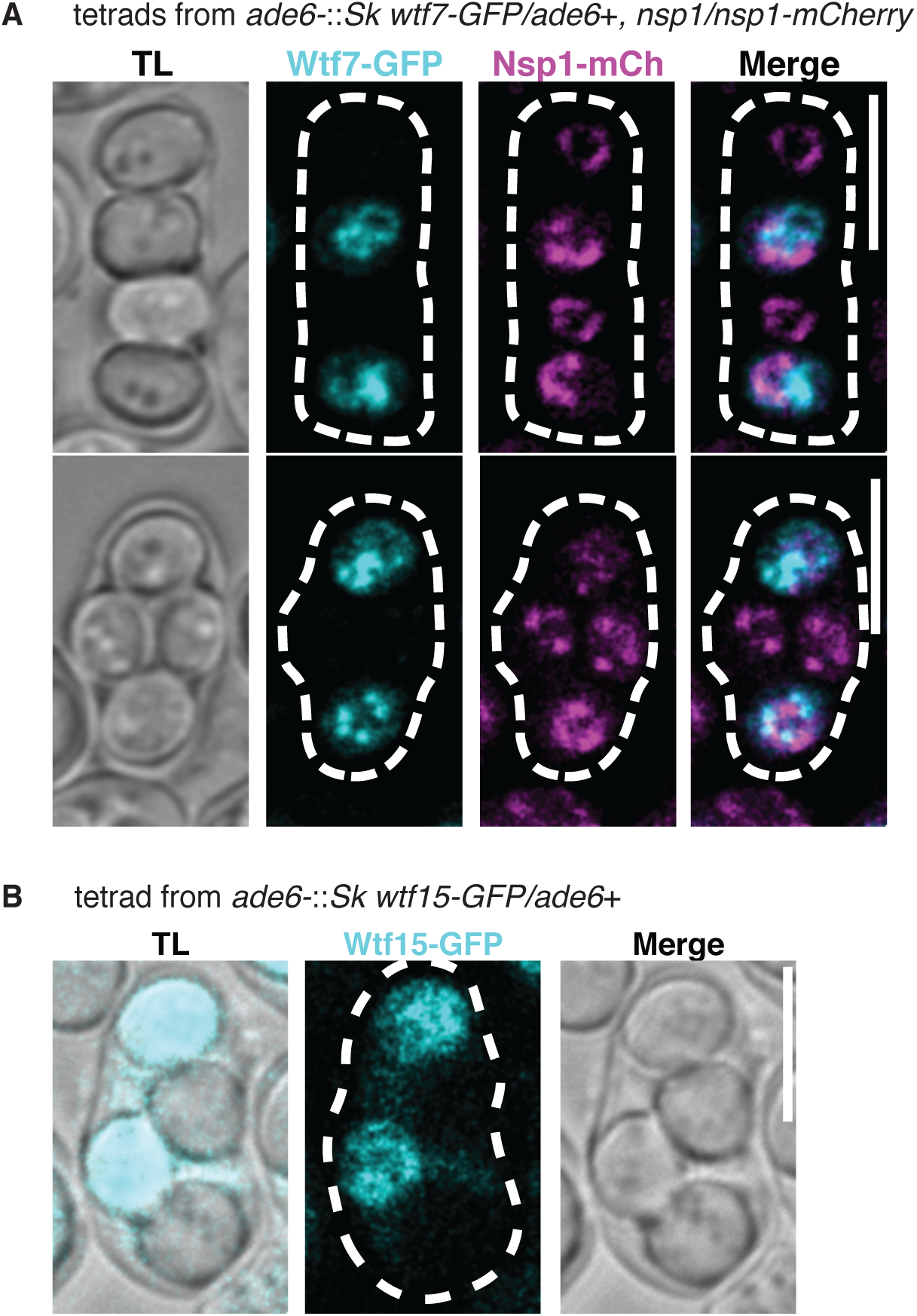
*Sk wtf7-GFP* and *Sk wtf15-GFP* are expressed in spores. (A) Representative images of *Sk* Wtf7-GFP localization in spores generated by diploids heterozygous for both *Sk wtf7-GFP* (green) and *nsp1-mCherry* (magenta; nucleoporin marker) are shown. (B) A tetrad generated from a diploid heterozygous for *Sk wtf15-GFP* is shown. We linearly unmixed these images (see Materials and Methods, Supplemental Figure 9). We adjusted the brightness and contrast differently for each image and smoothed them using Gaussian blur. Scale bar represents five microns. TL, transmitted light.

Unlike all other Wtf proteins, we saw expression of Wtf14 in vegetative cells. We saw that Wtf14-GFP colocalized with the endoplasmic reticulum (ER) marker *pbip1*::mCherry-AHDL in vegetative haploid and diploid cells (Figure 5A and 5B, Supplemental Figure 10) (28). This localization became more pronounced in cells during stationary phase (i.e. after ∼16 hours growth in rich media). These observations are consistent with previous work showing that *Sp* Wtf14-GFP expressed from an inducible promoter localizes to the ER (29) (Figure 5A and 5B, Supplemental Figure 10). We also saw ER localization of Wtf14-GFP in two of the four spores generated by diploids heterozygous for *wtf14-GFP* (Figure 5C). Similar to what has been observed in *Saccharomyces cerevisiae,* ER proteins seem to be generated *de novo* in spores (30). This localization pattern likely reflects expression of *wtf14-GFP* in the two spores that inherited the tagged *wtf14* allele.

## Discussion

### *S. pombe* genomes harbor a surprising number of active drive genes

Overall, we identified eight new *wtf* genes capable of meiotic drive. The *S. pombe* isolates with assembled *wtf* sequences contain between 4-14 predicted drivers (17, 21). Our current work shows that many of these predicted drive genes are able to drive in the right genetic context. This work expands the rapidly growing number of *bona fide* drive genes identified in recent years (4, 16–18, 31–41).

Moreover, our results highlight that one species can house a surprising number of functional meiotic drive genes. This is not unique to *S. pombe* as several species are known to harbor multiple gamete-killing drivers. These species include plants, fungi, insects, and animals (16–18, 25, 32–36, 39, 40, 42–45). For example, in *Drosophila simulans* there are three genetically distinct X-linked male drive systems. Two of these systems are paralogous, whereas the third arose independently (44, 46–48). Additionally, species can also house multiple copies of ‘true meiotic drive loci’ that bias chromosome segregation as well. Maize, for example, contains different numbers and sizes of heterochromatic regions (knobs) that have preferential segregation during female meiosis (49–51). Similarly, drive of more than one centromere has been observed in species of *Mus* (52, 53).

In *S. pombe*, one is compelled to ask how a genome can contain so many genes that act to destroy gametes. The costs of the selfish *wtf* genes could be at least partially offset because each genome also harbors multiple suppressors of drive. We speculate that we failed to observe drive of *Sp wtf23, Sk wtf27,* and *Sk wtf29* in this study due to the presence of suppressors. Consistent with this idea, we identified three new *wtf* genes in this work that act to suppress drive. We also found that some *wtf* genes can simultaneously promote their own drive while suppressing the drive of other *wtf* genes. Finally, our new findings reinforce the idea that Wtf antidotes must be highly similar to the poisons they neutralize (18). Similarity between drivers and suppressors has also been observed in other systems and this similarity could potentially serve as a general guide to identify suppressors (34, 35, 48).

In many ways, the landscape of meiotic drive genes in *S. pombe* is similar to that in *Podospora*. This filamentous fungus also contains multiple meiotic drive loci, including the *het-s* drive system in *P. anserina*, the best understood driver in any system at the molecular level (43, 54). In addition, *Podospora* contains its own multi-gene family of single-gene poison-antidote meiotic drivers, the *Spok* spore killers (34, 35). Like the *wtf* genes, the *Spok* genes are variable in number and location between isolates, leading to varying degrees of spore-killing phenotypes in crosses. Different isolates of *Podospora* contain various *Spok* genes (*Spok1, Spok2, Spok3,* and *Spok4*). Like *wtf* drivers, some *Spok* genes can both drive and suppress other drivers (34, 35). Similar to FY29033 *wtf18* and *Sp wtf13* in which suppression is bidirectional, *Spok1* and *Spok4* confer mutual resistance to each other (35). However, this case is not general among the *Spok* drivers. For example, the *Spok1* gene is resistant to the *Spok2* driver, but not *vice versa* (34).

The *wtf* genes also show some marked differences from the *Spok* gene family. Firstly, the same sequence appears to encode both the poison and antidote functions for a given *Spok* gene. In addition, the driving *wtf* genes are diverse in amino acid identity (30-90%), whereas the *Spok* genes are more conserved (>94% DNA sequence identity) within *P.* anserina. This could be due to the predicted enzymatic (nuclease and kinase) function of Spok proteins, which may constrain their divergence (34, 35).

Interestingly, although both *wtf* and *Spok* spore killers evolved independently and use different genes to enact drive, the parallels suggest that there could be recurrent themes shared by different drive systems. These characteristics could perhaps be exploited to aid in the discovery and characterization of novel natural drive systems, especially in organisms with low genetic tractability that show evidence of meiotic drive. Additionally, these shared themes could help us understand how artificial gene drive systems may evolve in populations.

### The potential functions of class 3 *wtf* genes remain largely unknown

Our final goal in this work was to test the potential functions of the class 3 *wtf* genes: *wtf7, wtf11, wtf14,* and *wtf15*. We observed no context in which we could detect Wtf11-GFP, but the proteomics data indicates it is expressed late in meiosis (27). We found that none of the class 3 *wtf* genes are required for vegetative growth. Additionally, our results showed that *wtf7, wtf11, wtf14,* and *wtf15* cannot drive at their endogenous loci, nor are they required to enable the drive of other *wtf* genes. However, we did find that *wtf7*, *wtf14,* and *wtf15* are expressed in a spore-specific manner, similar to the antidotes encoded by *wtf* drivers and suppressors. The functional significance of the class 3 *wtf* genes is unclear as none of these genes are required for fertility. Finally, we observed that Wtf14-GFP is localized to the ER during vegetative growth and in spores. This is the first observed expression of a Wtf protein during vegetative growth, demonstrating the potential for this gene to function outside of gametogenesis.

### Building a *wtf* meiotic driver

Our studies have elucidated that the ‘rules’ underlying the construction of *wtf* meiotic drivers are curiously lax. Firstly, a surprisingly wide breadth of sequences are equally capable of enacting drive. The sequence encoded in exon 1, found in antidote messages, is the most conserved (68-100% amino acid identity), suggesting this region may have specific interacting partners. The rest of the Wtf proteins is strikingly diverse. For example, the *Sk* Wtf9 and FY29033 Wtf35 proteins share <30% amino acid identity (Supplemental Figure 2B). However, both genes can drive into >90% of the functional gametes generated by a heterozygote.

In addition, our new analysis of the FY29033 *wtf35* driver revealed that the 7-amino acid repeat sequences encoded in the last exon of the characterized 6-exon *wtf* drivers are dispensable for drive. This is surprising because we previously found that a matching numbers of repeats in the Wtf^antidote^ and Wtf^poison^ proteins was important for poison-antidote specificity (18). The fact that FY29033 *wtf35* drives without these repeats suggests that they may function as spacers between more functionally important flanking domains. The important feature could be that having the same distance between the flanking domains helps to create compatible poison and antidote protein pairs.

Overall, the incredible diversity of *wtf* drivers could indicate that the toxicity of the Wtf poisons is not due to a specific enzymatic activity of the proteins or the targeting of a shared interactor. Rather, general shared features of Wtf proteins, such as the multiple predicted transmembrane domains, may be important.

The biggest constraint within *wtf* drive systems seems to be that antidotes must be very similar to the poisons they suppress. Within complete drivers, the absolute identity between the poison and antidote is guaranteed as they are both encoded on the same sequence. When an antidote is encoded by a locus distinct from the driver, it can still be effective at neutralizing a given poison. However, these sequences need to be highly similar. Although it is not yet clear exactly how much difference can be tolerated, the similarity at the C-terminus of the protein is particularly important. All known antidotes are >96% similar in amino acid sequence to the poisons they neutralize. Future studies of the mechanisms used by the Wtf proteins will be guided by and hopefully reveal the molecular basis of these features.

## Materials and Methods

### Viable spore yield assay and allele transmission

To assay fertility and allele transmission, we began by generating heterozygous diploids as described in (25). We grew the haploid parental strains in YEL (0.5% yeast extract, 3% dextrose, and 250 mg/L adenine, histidine, leucine, lysine, and uracil) to saturation at 32°C. Using the saturated cultures, we added 300 microliters of each strain to an Eppendorf tube and vortexed it to mix the cells. We spun these cells down and resuspended the cell pellet in sterile ddH_2_O. We then plated 100 microliters of the cell mixture onto SPA (1% dextrose, 7.3mM KH_2_PO_4_, vitamins, and agar) or SPAS (SPA + 45mg/L adenine, histidine, leucine, lysine, and uracil) plates and incubated the plates at 25°C for ∼16 hours. This allowed the haploids to mate and form diploids. Because some of the strains we used are homothallic (meaning they can switch mating type), we selected for heterozygous diploids. Most of our haploid parental strains had complementary auxotrophic markers which allowed us to select for heterozygous diploids by streaking the mated cell mix on minimal media. We then restreaked colonies that grew on minimal media to further isolate single colonies on minimal media. However, some of the heterozygous diploids were adenine auxotrophs. To select those diploids, we did the same procedure using minimal media supplemented with 45 mg/L of adenine.

Next, we cultured the heterozygous diploids in YEL overnight to saturation in a 32°C shaker. We then plated 50 microliters of our saturated culture onto SPA to allow the diploids to sporulate (for three days at 25°C). We also diluted the cultures and subsequently plated the cells onto YEA (YEL + agar), and let them grow for three days at 32°C. These plates were then replica-plated to minimal media (+ adenine for adenine auxotrophs) as well as any other selective media to further confirm we isolated heterozygous diploids. From the YEA plates, we also calculated the colony-forming units (CFU) to determine the concentration of viable diploids in the YEL culture. We next scraped the cells off of the three day old SPA plates into 500 microliters of sterile ddH_2_O and treated them with five microliters of glusulase (Sigma-Aldrich) for four hours at 32°C to shed the ascal membrane and wall to release the spores (55). We then killed any remaining vegetative cells by adding 500 microliters of 60% ethanol for 10 minutes at room temperature. Next, we washed the spores in ddH_2_O and resuspended them in 500 microliters of sterile ddH_2_O. We diluted the spores and plated them onto YEA and let them grow into colonies at 32°C for three to five days to determine the CFU. Additionally, we picked colonies from the YEA plate onto a YEA master plate and grew the plate at 32°C for ∼24 hours. We then replica-plated the master plate to yeast nitrogen-based plates with a specific dropout of either adenine, histidine, uracil, lysine, leucine, and plates containing various drugs (G418, Hygromycin B, and Nourseothricin) to determine the genotype of each spore and thus assay allele transmission.

### Viable spore yield calculations

To determine the fertility of selected stable heterozygous diploids, we calculated the viable spore yield (number of viable spores recovered from SPA/ the number of viable diploid cells plated on SPA) (55). The number of viable cells used in these calculations was determined using the CFU counts described above.

To determine the viable spore yield of homothallic haploids (i.e. SZY2254 and SZY3529), we first grew the strains in YEL cultures overnight at 32°C with shaking. We then spread 100 microliters of each saturated culture onto SPA plates and left the plates at 25°C for three days. This step allowed the haploid cells to mate and subsequently sporulate. Using the same culture, we also performed serial dilutions and plated them onto YEA to quantify the number of haploids in the original culture. After three days, we scraped the spores off of the SPA plates and treated them as described above. Next, we performed a series of dilutions and plated the spores onto YEA plates to quantify the number of viable spores. We then incubated these plates at 32°C for five days. To determine the fertility, we calculated the viable spore yield (number of viable spores recovered from SPA/ number of haploids plated on SPA).

### Strain construction: *ade6-*integrating vectors

All strain names and genotypes are presented in Supplemental Table 4. To assay allele transmission of different *wtf* genes, we used *ade6-*integrating vectors containing *kanMX4* or *hphMX6* drug resistance markers (56, 57). pSZB188 (empty vector with *kanMX4* resistance) was published in (16). pSZB386 (empty vector with *hphMX6* resistance) was published in (18). pSZB387 is identical to pSZB386. These vectors have a KpnI site within a mutant *ade6-*targeting cassette that we cut to linearize the plasmid. We then introduced plasmids into yeast using a standard lithium acetate protocol. Proper integration of these vectors at *ade6* yields an Ade*-* phenotype (red colonies). To make the *ade6-*integrating vectors with the *wtf* transgenes, we cloned the *wtf* alleles using the oligos, DNA templates, restriction enzymes, and target sites described in Supplemental Table 5.

We could not directly amplify FY29033 *wtf1* alone from the genome, due to repetitive nature of the *wtf* genes. Instead, we first amplified both FY29033 *wtf1* + *wtf36* tandem genes together from the FY29033 strain using oligos 1346 and 1348. From that PCR product, we amplified the FY29033 *wtf1* allele using oligos 1352 and 1592. We then digested this fragment with SacI and subsequently ligated this allele into the SacI site of pSZB387 to make pSZB879. The plasmids are described in Supplemental Table 6.

### Deleting Sk wtf1, Sp wtf7, Sp wtf11, Sp wtf14 + wtf15

To generate an *Sk wtf1* deletion cassette, we first amplified the regions (∼750bp) upstream and downstream of the *wtf1* locus using oligo pairs 645+656 and 2158+646, respectively. The upstream region includes the Tf1 transposon flanking the left side of the *Sk wtf1* locus. For these PCR reactions, we used SZY661 as a template. We then amplified the *hphMX6* casette from pAG32 with oligos 657 and 2159 (57). We stitched all three fragments together using overlap PCR and transformed this deletion cassette into SZY298 to generate SZY3829. We confirmed the deletion using a series of PCR reactions: two oligo pairs with one oligo external to and one oligo within the deletion cassette (660+AO638 and AO1112+661) and a pair of oligos with one oligo outside of the deletion cassette and one oligo internal to the *Sk wtf1* locus (660+2287).

To generate an *Sp wtf7* deletion cassette, we first began by amplifying the regions (∼1 kb) upstream and downstream of the gene with oligos 1565 + 1566 and oligos 1569 + 1570, respectively. We then amplified the drug cassettes, either *hphMX6* or *natMX4*, with oligos 1567 and 1568 using pAG32 or pAG25 as templates, respectively (57). These oligos contained tails with homology to the upstream and downstream sequence of *Sp wtf7*. We then stitched these separate fragments together using overlap PCR and subsequently transformed it into yeast using the standard lithium acetate protocol. We deleted *Sp wtf7* in the SZY44 strain to generate SZY2309, and deleting *Sp wtf7* in SZY643 generated strains SZY2310 and SZY2336. We confirmed these deletions using a series of PCR reactions: oligo pairs with one oligo outside of the deletion cassette and one oligo internal to the gene (1571+1586 and 1572+1585), and 2 oligo pairs in which one oligo was external to and one oligo was within the deletion cassette (1571+AO638 and 1572+AO1112). Additionally, we further confirmed deletion of the gene using oligos internal to *Sp wtf7* (2152 and 2153).

To generate an *Sp wtf11* deletion cassette, we began by amplifying the regions (∼500 bp) upstream and downstream of the gene using oligos 1667+1669 and oligos 1670+1672, respectively. Next, we amplified the *hphMX6* drug cassette with oligos 1668 and 1671 using pAG32 as a template (57). We put these fragments together using overlap PCR and transformed it into SZY2336 to generate strains SZY2854 and SZY2855. This deletion was confirmed with two pairs of oligos with one oligo outside of the deletion cassette and one oligo internal to *wtf11* (1705+1707 and 1706+1707) and 2 pairs of oligos in which one oligo was external to the deletion cassette and one oligo was internal to the *hphMX6* cassette (1705+AO638 and 1706+1842). We further confirmed the deletion of the gene with a PCR reaction using oligos internal to *Sp wtf11* (2154 and 2155).

To delete the tandem *Sp wtf14* and *wtf15* genes, we first made a deletion cassette via PCR. To do this, we amplified the upstream region (∼600 bp) of *Sp wtf14* using oligos 1649 and 1651, and the downstream region (∼1 kb) of *Sp wtf15* with oligos 1655 and 1657. We also amplified the *natMX4* drug cassette from pAG25 with oligos 1650 and 1656 and then stitched the three fragments together using overlap PCR (57). We then transformed this construct into SZY643 to generate strains SZY2856 and SZY2857. We confirmed the deletion of *Sp wtf14* and *Sp wtf15* via PCR using two oligo pairs in which one oligo was external to the deletion cassette and one oligo was within the *wtf14* and *wtf15* locus (1709+1710 and 1658+1711). We also performed a PCR with in which one oligo was external to the deletion cassette and one was internal to the *natMX4* cassette (1709+AO638 and 1658+1843). Additionally, we did a PCR with oligos within the *Sp wtf14 + wtf15* locus (2156 and 2157) to confirm the absence of the genes.

To assay the effect of the class 3 *wtf* genes, we generated strains lacking *wtf7, wtf11, wtf14,* and *wtf15* genes in *Sp.* First, we exchanged the *natMX4* gene marking the *wtf14+wtf15* deletion with the *CaURA3MX* cassette as described in (58). Briefly, we amplified the *CaURA3MX* cassette from pFA6-mTurq2-URA3MX using oligos PR78 and PR79 (59). We then transformed this fragment into SZY2856 to generate the yeast strain SZY3448. We then generated the quadruple deletion mutant strain via crosses.

### Spot assay

To determine if a mutant strain lacking class 3 *wtf* genes had growth defects, we first cultured the strains in five ml of YEL at 32°C with shaking overnight. We then diluted the cultures to an OD600 of 0.1 and grew them for six hours at 32°C with shaking. We then did serial dilutions and spotted five microliters onto YEA plates and incubated them at 32°C for 2 days.

### Deleting *Sp wtf21* using CRISPR

We used the *S. pombe* CRISPR-Cas9 genome editing system from (60) to generate the *Sp wtf21*Δ::*kanMX4* mutation in SZY890. This system uses two plasmids. The first one expresses a guide RNA to the target sequence and the second plasmid expresses Cas9 (pMZ222). To generate the plasmid carrying a guide RNA targeting *Sp wtf21* (pSZB197), we annealed oligos 623 and 624 to each other and ligated them into the CspCI site of pMZ283. Next, we transformed pMZ222 and pSZB197 into *Sp* (SZY643) along with a *wtf21*Δ::*kanMX4* repair cassette (see below). We initially selected for yeast that contained both plasmids (Leu+ Ura+) and subsequently selected for G418-resistant colonies. We then screened for the desired mutants by PCR using oligos 631 and 632 that flank *wtf21*, but are outside of the region amplified in the repair cassette. We generated the repair cassette using PCR to build a fragment containing the regions upstream and downstream of *wtf21* flanking the *kanMX4* gene (56). We amplified the upstream and downstream regions with oligos 625+626 and 629+630, respectively, using *Sp* genomic DNA as a template. We amplified the *kanMX4* gene with oligos 627+628. We then stitched the three fragments together using overlap PCR to generate the repair cassette.

### C-terminally GFP-tagged *wtf* alleles

We generated *Sk wtf7, wtf11,* and *wtf15* GFP-tagged alleles using the following strategy. We first amplified the *Sk wtf* alleles (*wtf7, wtf11,* and *wtf15*) with their endogenous promoters using genomic DNA from SZY661 using oligo pairs 1359+1360 for *wtf7*, 1362+2210 for *wtf11,* and 991+1368 for *wtf15.* We then amplified yEGFP using pKT127 (61) as a template with oligos 1361+634 for *wtf7,* 2211+634 for *wtf11,* and 1369+634 for *wtf15.* We then stitched the two respective fragments together using overlap PCR and digested the fragments with SacI. We then cloned these fragments into the SacI site of pSZB188 to generate pSZB691 (*wtf7-GFP*), pSZB1087 (*wtf11-GFP*), and pSZB698 *(wtf15-GFP)*.

To generate *Sk wtf14-GFP,* we amplified *wtf14* from pSZB378 using oligos 1365 and 1366. To amplify yEGFP, we used oligos 1367 and 634 and used pKT127 as a template (61). We then used overlap PCR to stitch the two fragments together and digested the fragment with SacI. We then cloned it into the SacI site of pSZB188 to generate pSZB696.

### Imaging GFP-tagged Wtf proteins

To determine the localization of Wtf7-GFP, Wtf11-GFP, Wtf14-GFP, and Wtf15-GFP during haploid and diploid vegetative growth, we imaged cells during logarithmic and stationary phase. To image the cells during stationary phase, we first made a five ml YEL culture of each haploid and diploid strain and grew them for 16 hours in a 32°C shaker. We then imaged these cells as our stationary phase samples. To obtain cells in logarithmic phase, we used the YEL cultures described above and diluted them 1:10 in five ml cultures and incubated the tubes with shaking at 32°C for four hours. To image the cells, we spotted 3 microliters of culture onto a glass slide pre-coated with lectin and covered with a glass coverslip to keep the cells in place for imaging.

To image the spore sacs, we used diploids that had sporulated on SPA plates at 25°C for two days. To prepare cells for imaging, we first scraped the cells off of SPA plates and mixed them with 3 microliters of lectin. We then plated them on glass slides and covered with a glass coverslip.

For all experiments, we imaged the cells on an LSM-780 (Zeiss) AxioObserver confocal microscope with a 40X C-Apochromat water-immersion objective (Zeiss, NA= 1.2) or a 40X LD C-Apochromat water-immersion objective (Zeiss, NA=1.1). We acquired images of every field of cells in two ways. We acquired a channel mode image to obtain a transmitted light image. For the images in Figure 5C, we acquired the fluorescence images by exciting GFP at 488 nm and collecting its emission between a 500-553 bandpass filter, while we excited mCherry at 561 nm and collected its emission between a 562-615 nm bandpass filter. For all other images, we acquired images in lambda mode over the entire visible range, with GFP and mCherry excitation at 488 and 561 nm, respectively, to obtain the true fluorescence. To eliminate cross talk, we collected the GFP and mCherry lambda images separately. To distinguish true GFP and mCherry signal from autofluorescence, we linearly unmixed the lambda mode data for GFP and mCherry using reference GFP and mCherry images and an in-house plug-in in ImageJ (https://imagej.nih.gov/ij/).

### Analysis of meiotic proteomic data

We used the data set collected by Krapp et al and determined the relative protein levels following the method described in (27). We considered all Wtf proteins that were detected in at least one timepoint in at least one of the three replicate experiments. We plotted the mean of the quantified Wtf protein for each timepoint that had at least two replicates, regardless of the number of replicates at that specific timepoint. When a Wtf protein was detected in only a single replicate, we plotted that value for that timepoint.

### Alignments

To determine the DNA and amino acid sequence identity shared by *wtf* genes and proteins, we aligned the sequences using Geneious Prime (https://www.geneious.com). We used the Geneious alignment tool and performed a global alignment with free end gap using the default parameters. For DNA sequence alignments, the parameters we used were: cost matrix= 65% similarity, gap open penalty=12, and gap extension penalty=3. For protein sequence alignments we used Blosum62 as the matrix, gap open penalty=12, gap extension penalty=3, and refinement iterations=2.

## Acknowledgments

We thank Dr. Sue Jaspersen and members of the Zanders lab for their helpful comments on the manuscript. We are grateful to Joe Varberg and Sue Jaspersen for making and sharing their unpublished *nsp1-mCherry* strain, and Risa Mori and Snezhana Oliferenko for the *pbip1*-*mCherry* ER marker strain. This work was performed to fulfill, in part, requirements for MABN’s thesis research in the Graduate School of the Stowers Institute. Original data underlying this manuscript can be accessed from the Stowers Original Data Repository at http://www.stowers.org/research/publications/libpb-1433. This work was supported by the following awards to SEZ: The Stowers Institute for Medical Research (https://www.stowers.org), the March of Dimes Foundation Basil O’Connor Starter Scholar Research Award No. 5-FY18-58 (https://www.marchofdimes.org), the Searle Scholar Award, and the National Institutes of Health (NIH) under the award numbers R00GM114436 and DP2GM132936 (https://www.nih.gov). MABN was also supported by the National Cancer Institute of the NIH under award number F99CA234523. The funders had no role in study design, data collection and analysis, or manuscript preparation.

---

**Supplemental Figure 1.**
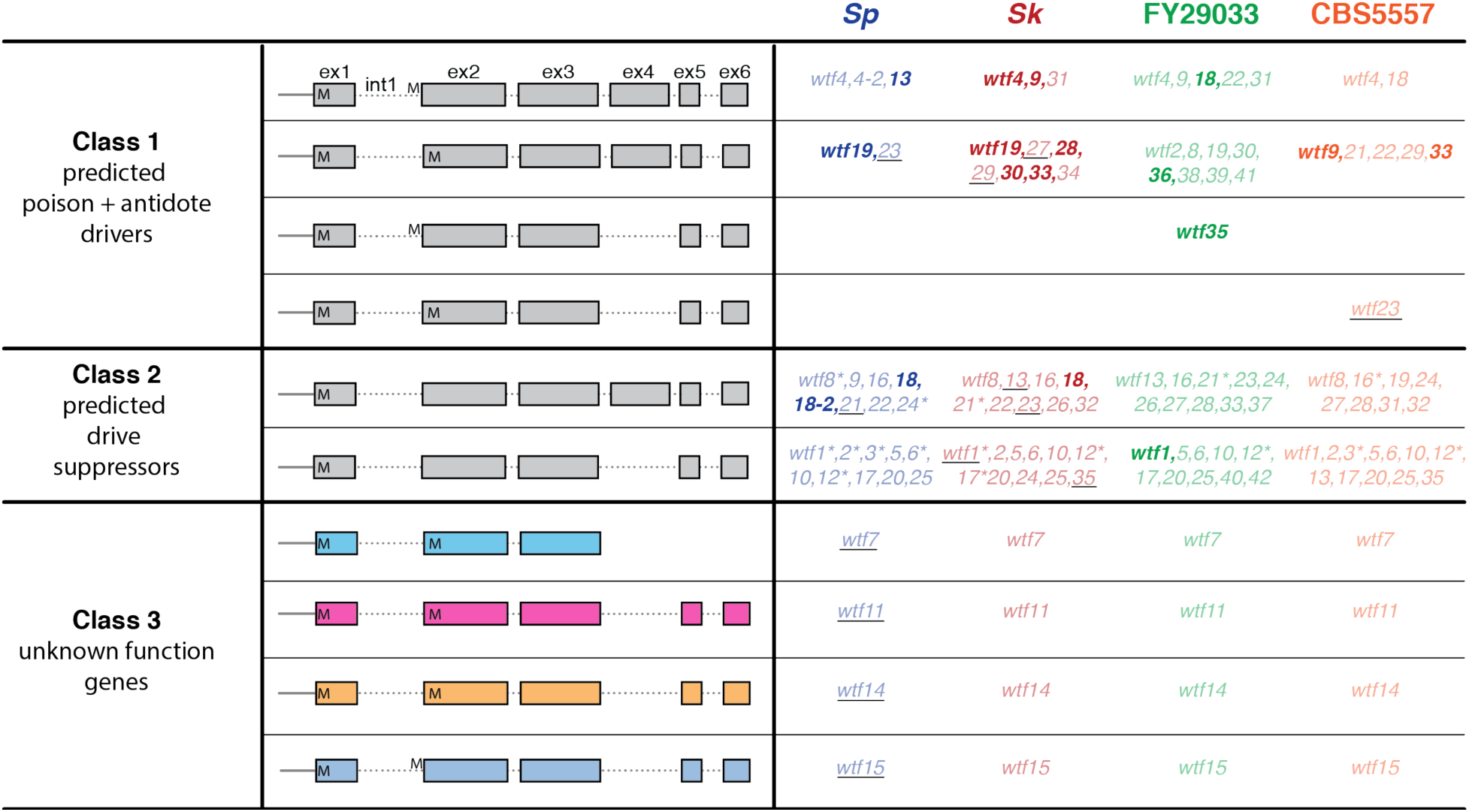
Classification of the *wtf* gene family based on their sequence and structure (21). Class 1 is predicted to contain poison-antidote meiotic drivers. Class 2 *wtf* genes are predicted to be antidote-only genes and suppressors of drive. *wtf7, wtf11, wtf14,* and *wtf15* are all grouped in class 3 solely because they do not look like any other *wtf* gene in this family, or each other. The *wtf* genes that have been shown in this study or were previously characterized as meiotic drivers or suppressors are shown in bold (16–18). The *wtf* genes that were tested in this study but did not show a meiotic drive phenotype are underlined. *wtf* genes with in-frame stop codons are depicted with a “*” next to the gene name and ‘M’ highlights the start codons and the in-frame ATG codons near the start of exon 2.

**Supplemental Figure 2.**
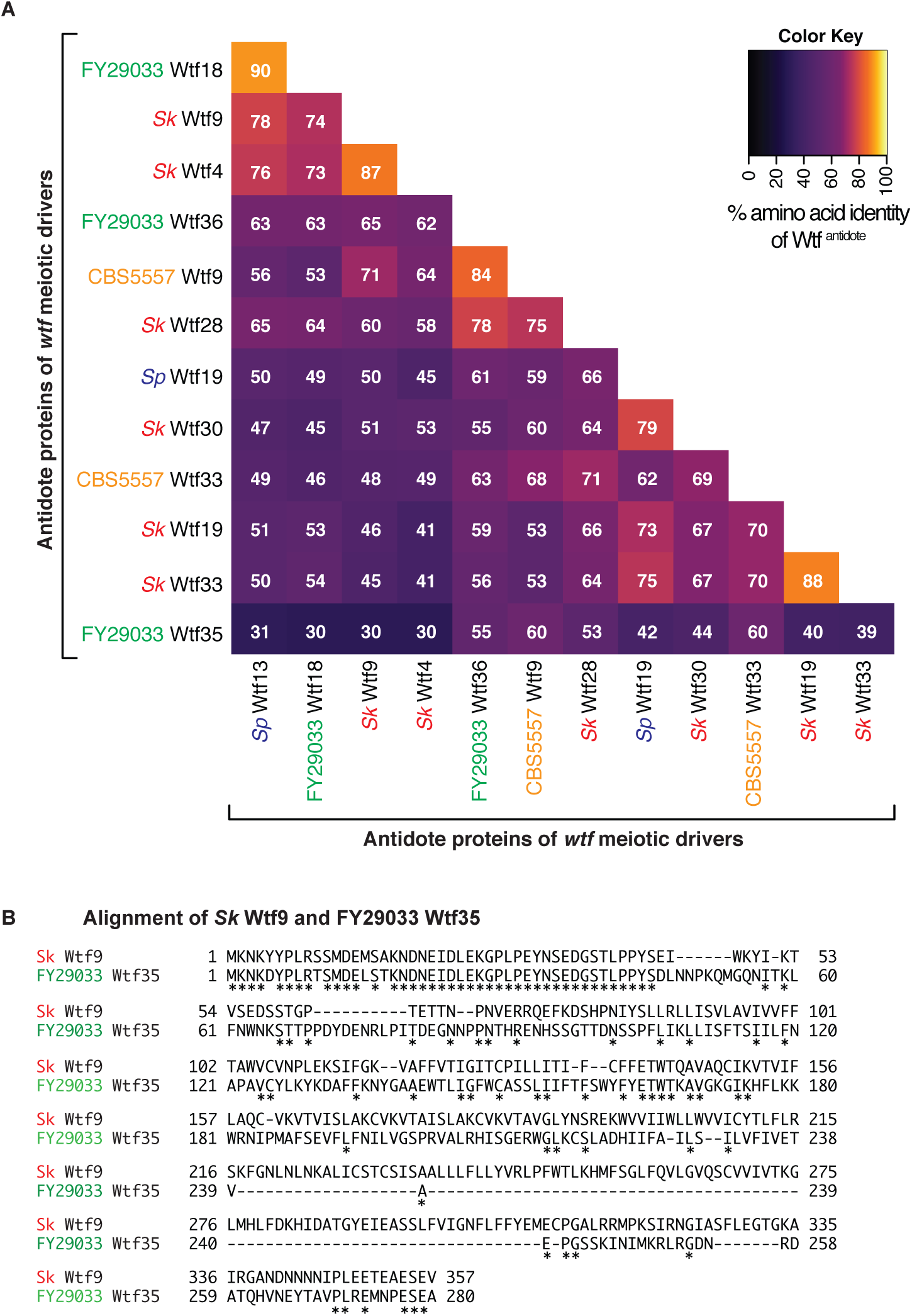
w*t*f meiotic drivers are extremely diverse. (A) The amino acid identity shared by pairs of Wtf^antidote^ proteins encoded by *bona fide* meiotic drivers are shown (16–18). (B) Manual alignment of the long isoforms of *Sk* Wtf9 and FY29033 Wtf35. Using this method, the percent identity between these two proteins is 24%.

**Supplemental Figure 3.**
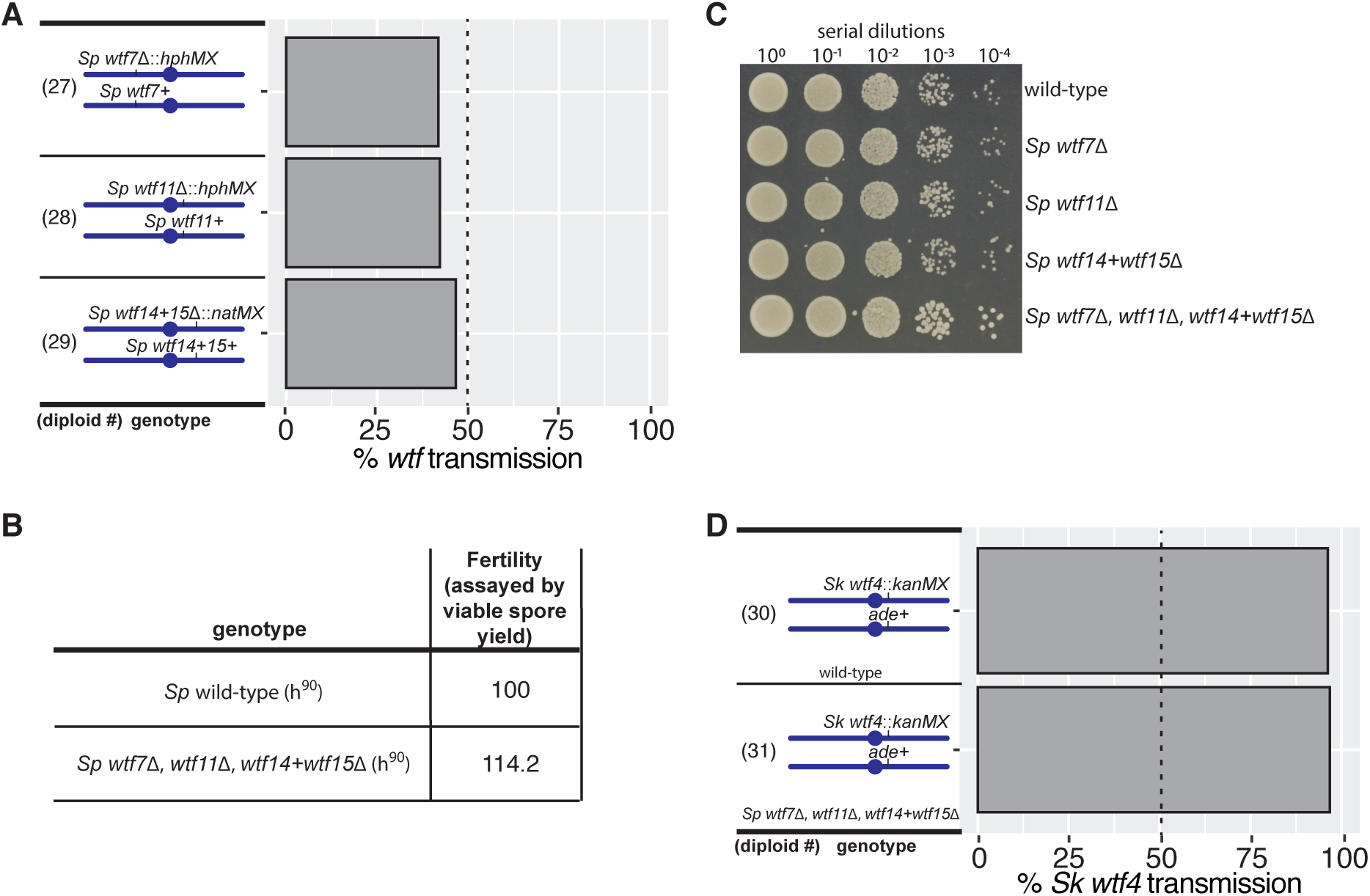
Class 3 *wtf* genes do not exhibit meiotic drive phenotypes. (A) Allele transmission into spores from diploids heterozygous for class 3 *wtf* genes is shown. No significant differences in the allele transmission were found using a G-test when compared to the control *ura4* locus. (B) Fertility of wild-type and mutant backgrounds (normalized to wild-type). No significant differences in the viable spore yield were found using a Wilcoxon test. The complete raw data are presented in Supplemental Table 2. (C) Serial dilutions of strains with the denoted genotype were spotted onto YEA plates. The slight growth advantage of the quadruple mutant is likely due to the fact that the mutant is a uracil prototroph while the others are uracil auxotrophs. All other auxotrophies are matched between the strains. (D) Allele transmission of the *Sk wtf4* meiotic driver in a wild-type and mutant background is shown. We found no significant difference between diploid 30 and diploid 31 using a G-test. For (A) and (D), we genotyped more than 200 spore progeny for each diploid and the complete raw data are presented in Supplemental Table 1.

**Supplemental Figure 4.**
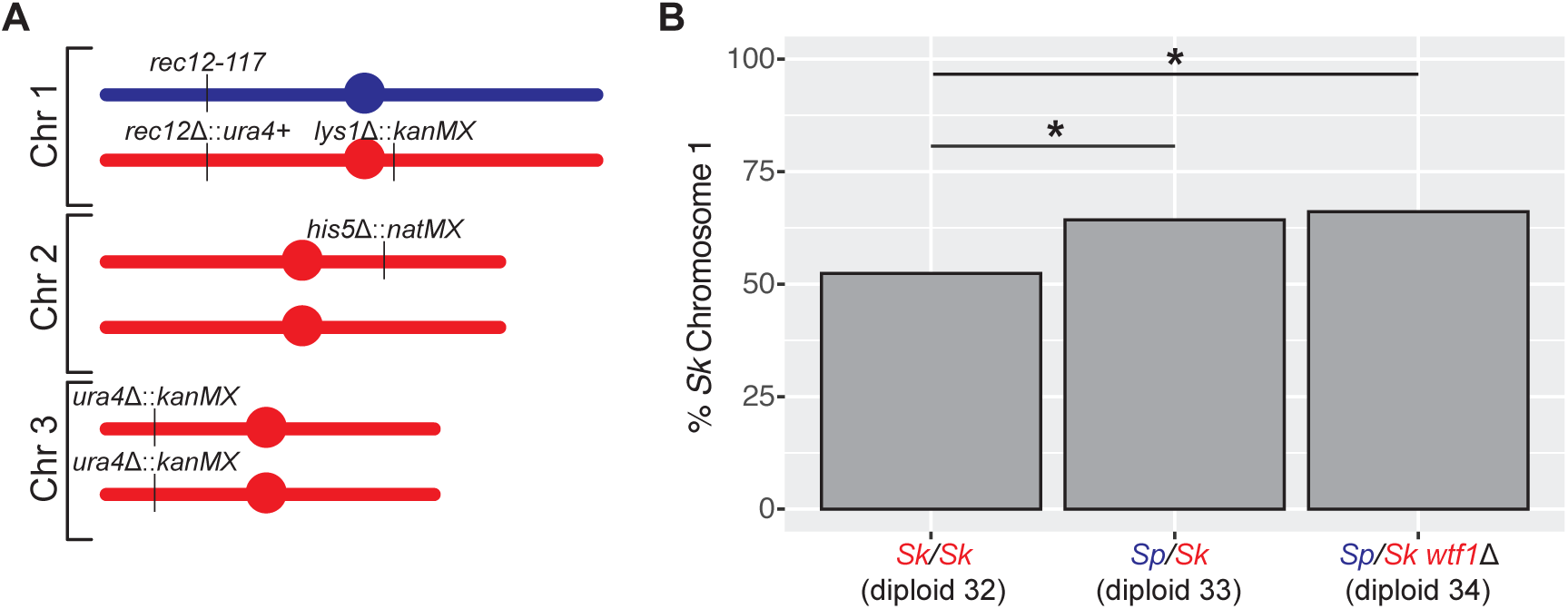
The allele transmission bias favoring *Sk* chromosome 1 is not caused by *wtf* genes. (A) Cartoon summary of the genotype of the *Sp/Sk* hybrid diploid. (B) Allele transmission of *Sk* chromosome 1 in *rec12-* diploids. We genotyped more than 400 haploid spore progeny from each diploid. * indicates a p-value<0.05 (G-test). The complete raw data is presented in Supplemental Table 3.

**Supplemental Figure 5.**
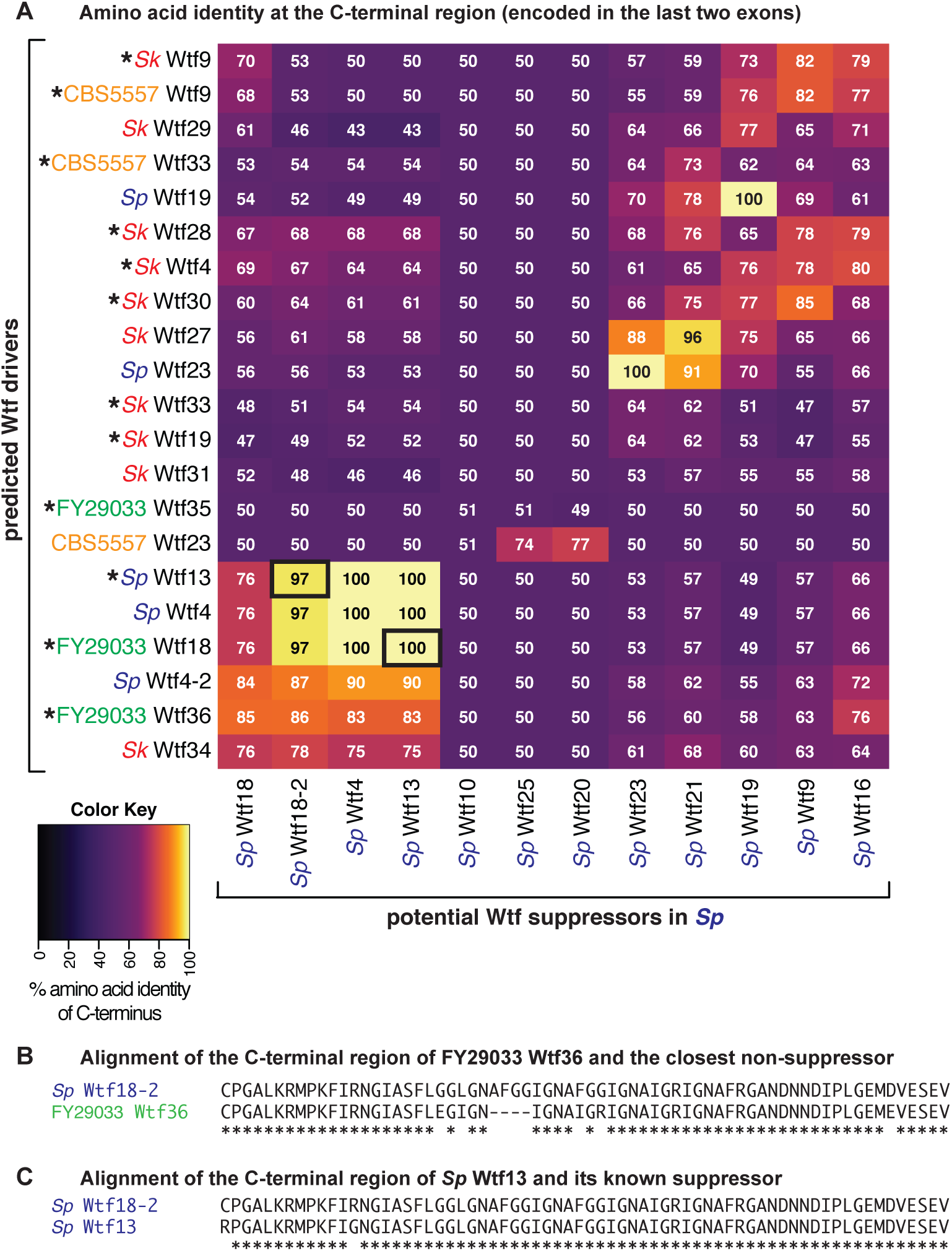
High amino acid identity appears to be required for suppression of *wtf* meiotic drive. (A) The amino acid identity of the C-terminal region (encoded in the last two exons) of pairs of Wtf^antidote^ proteins are shown. *wtf* meiotic drivers known to drive in an *Sp* background are depicted with a “*”. Characterized *wtf* driver-suppressor pairs are shown with black boxes. *wtf* genes were only considered as potential suppressors if they are transcribed during meiosis (22). (B) An amino acid sequence alignment of the C-terminal region of the FY29033 Wtf36 driver and *Sp* Wtf18-2, a highly similar *wtf* gene that does not suppress FY29033 *wtf36* is shown. (C) An amino acid sequence alignment of the C-terminal region of *Sp* Wtf13 and its suppressor *Sp* Wtf18-2 is shown (18).

**Supplemental Figure 6.**
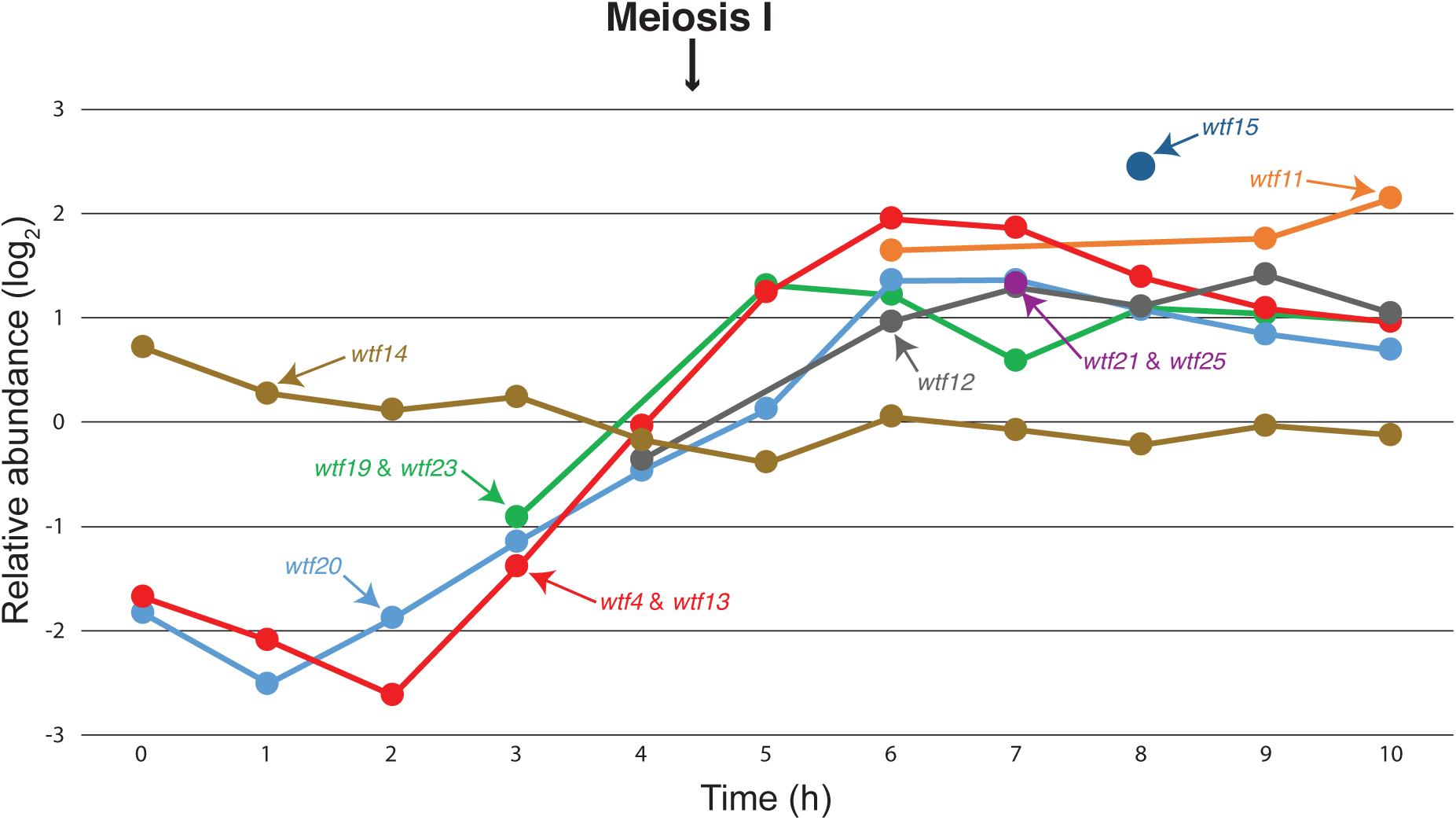
Class 3 Wtf proteins are present during meiosis. Analysis of Wtf protein levels during meiosis from the data of (27). Due to the high sequence similarity between *Sp* Wtf4 and Wtf13 proteins, the data points for these two proteins were merged.

**Supplemental Figure 7.**
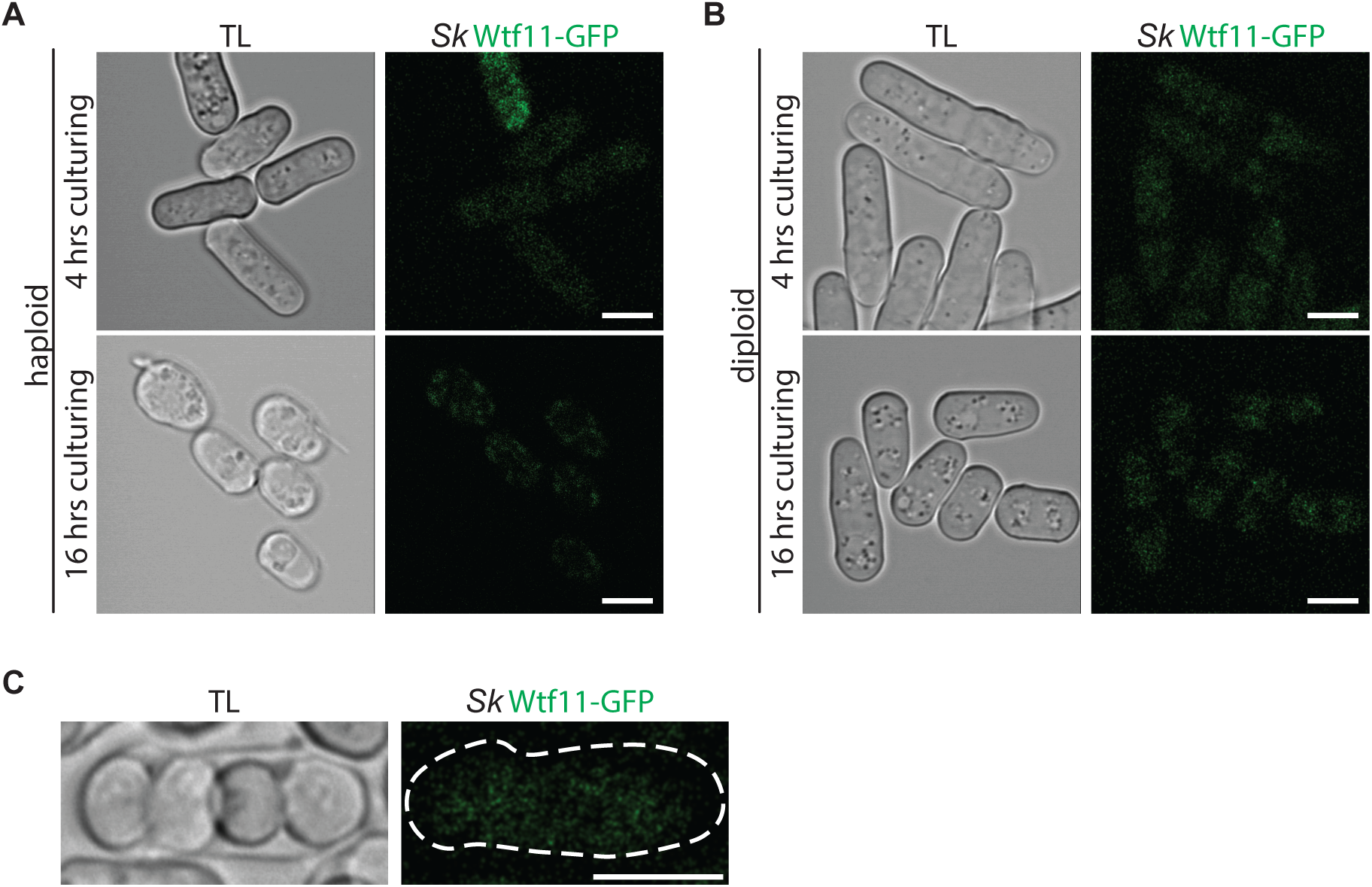
We failed to observe *Sk wtf11-GFP* expression during vegetative growth or sporulation. Representative images of cells containing *Sk* Wtf11-GFP (green) in (A) haploids, (B) heterozygous (*wtf11-GFP/ade+*) diploids, and (C) a tetrad generated by heterozygous diploids. We adjusted the brightness and contrast to observe the background and adjusted them differently for each image. We smoothed the images using Gaussian Blur. TL, transmitted light. Scale bar represents five microns.

**Supplemental Figure 8.**
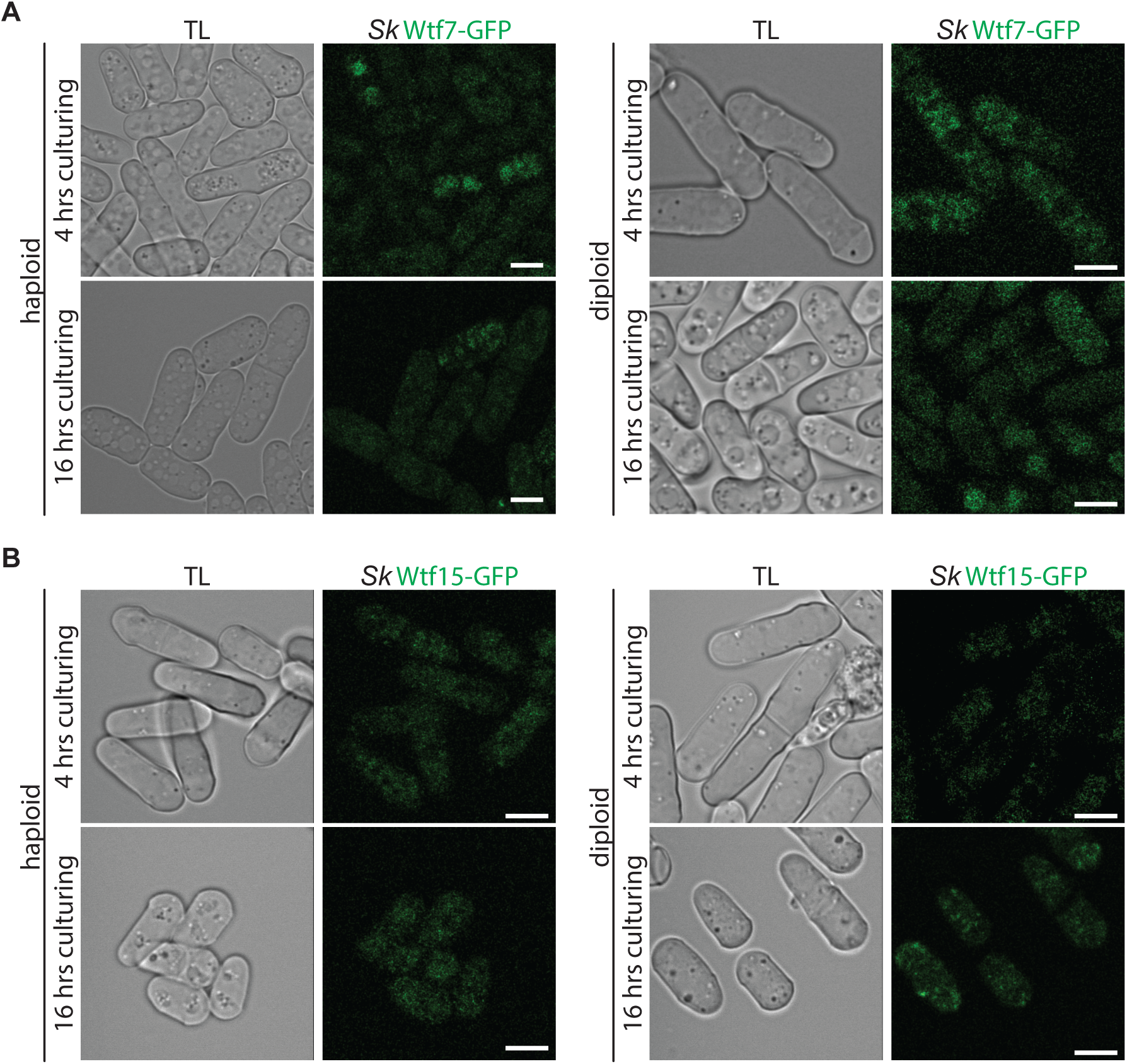
We did not observe *Sk wtf7-GFP* or *Sk wtf15-GFP* expression in vegetative haploids or diploids. (A) Representative images of Wtf7-GFP (in green) in haploids (left) and heterozygous diploids (right) during logarithmic phase and saturation are shown. (B) Representative images of Wtf15-GFP (in green) in haploids (left) and heterozygous diploids (right) during logarithmic phase and saturation are shown. We linearly unmixed these images and adjusted the brightness and contrast differently for each image and smoothed them using Gaussian Blur. The brightness and contrast were adjusted to observe the background. We verified the green autofluorescence via spectral imaging. The scale bar represents five microns. TL represents transmitted light.

**Supplemental Figure 9.**
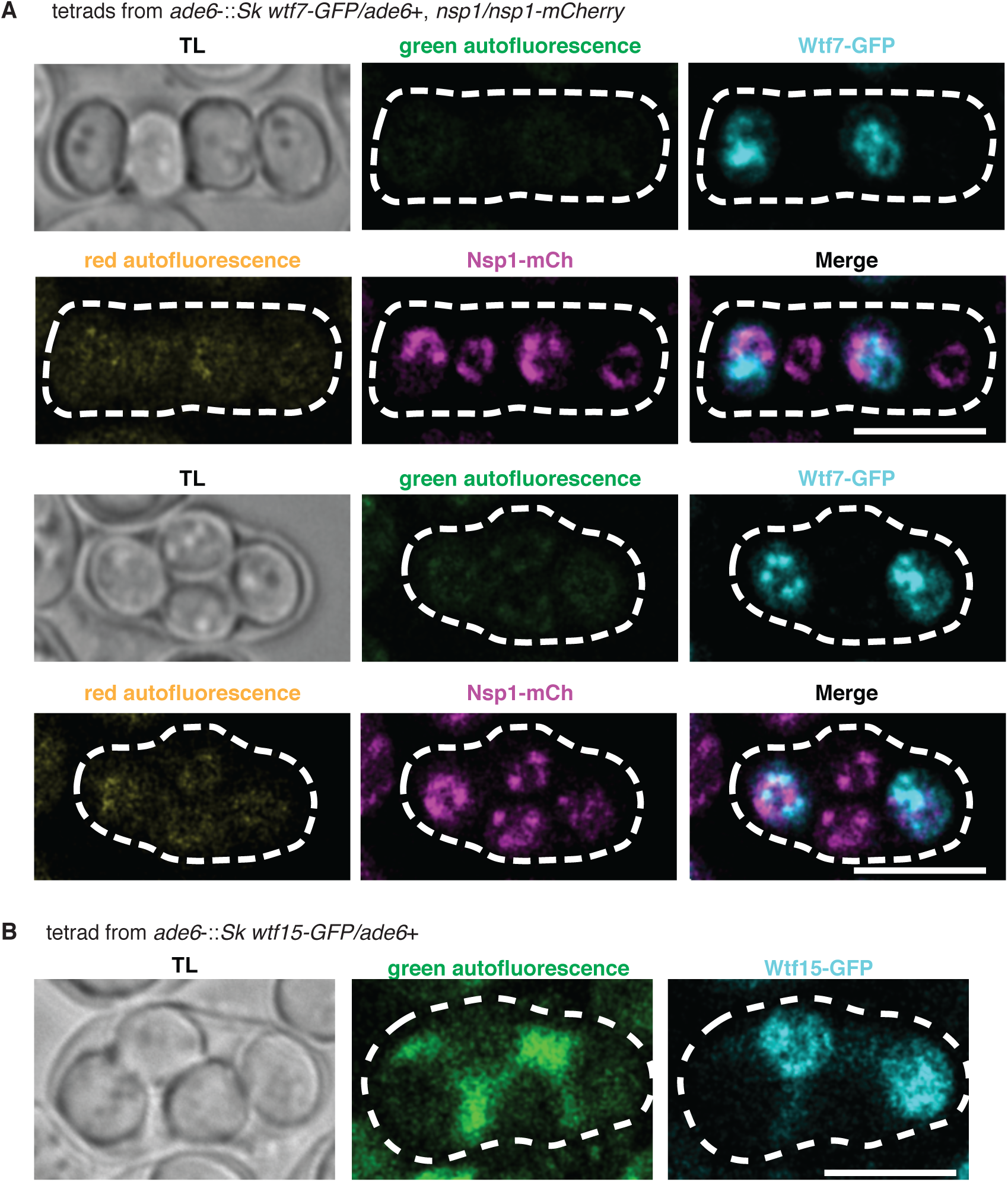
Linear unmixing of *Sk* Wtf7-GFP and *Sk* Wtf15-GFP localization in spores. (A) Linear unmixing of the representative images shown in Figure 4A of *Sk* Wtf7-GFP (cyan) and Nsp1-mCherry (magenta) is shown. We normalized the intensities of the green and red autofluorescence to the intensities of the GFP and mCherry signals, respectively. (B) Linear unmixing of the representative image shown in Figure 4B of *Sk* Wtf15-GFP (cyan) localization is shown. We normalized the intensity of the green autofluorescence to the intensity of the GFP channel. We adjusted the brightness and contrast differently for each image and smoothed them using Gaussian blur. The scale bar represents five microns and TL represents transmitted light.

**Supplemental Figure 10.**
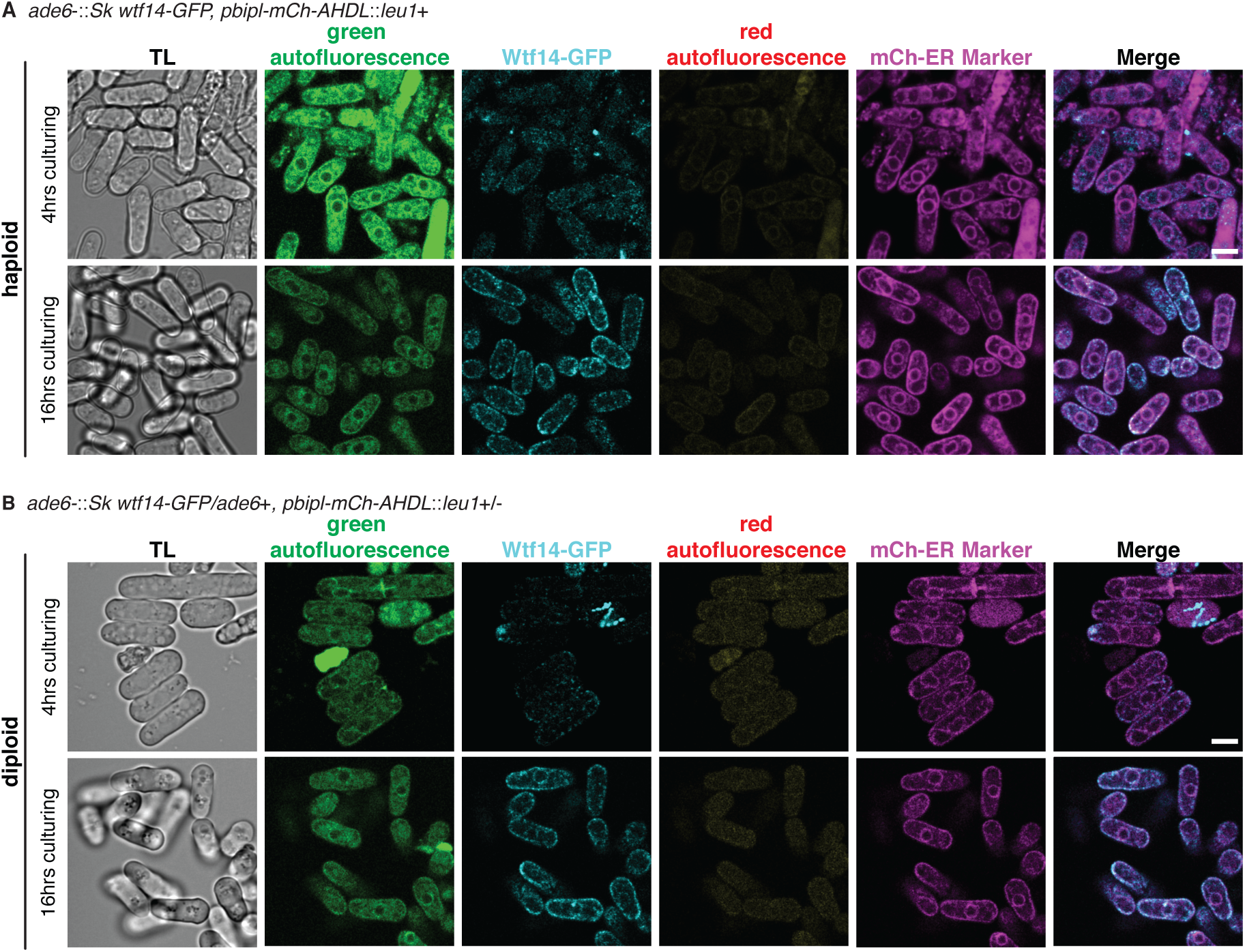
Linear unmixing of *Sk* Wtf14-GFP in haploid and diploid cells. Linear unmixing of (A) haploids and (B) diploids containing *Sk* Wtf14-GFP (cyan) and mCherry-AHDL (magenta) from the representative images shown in Figure 5A and 5B (28). We adjusted the intensities of the autofluorescence images to their respective channels. We adjusted brightness and contrast differently for each image and smoothed them using Gaussian blur. TL represents transmitted light. Scale bar represents five microns.

**Supplemental Table 1.**
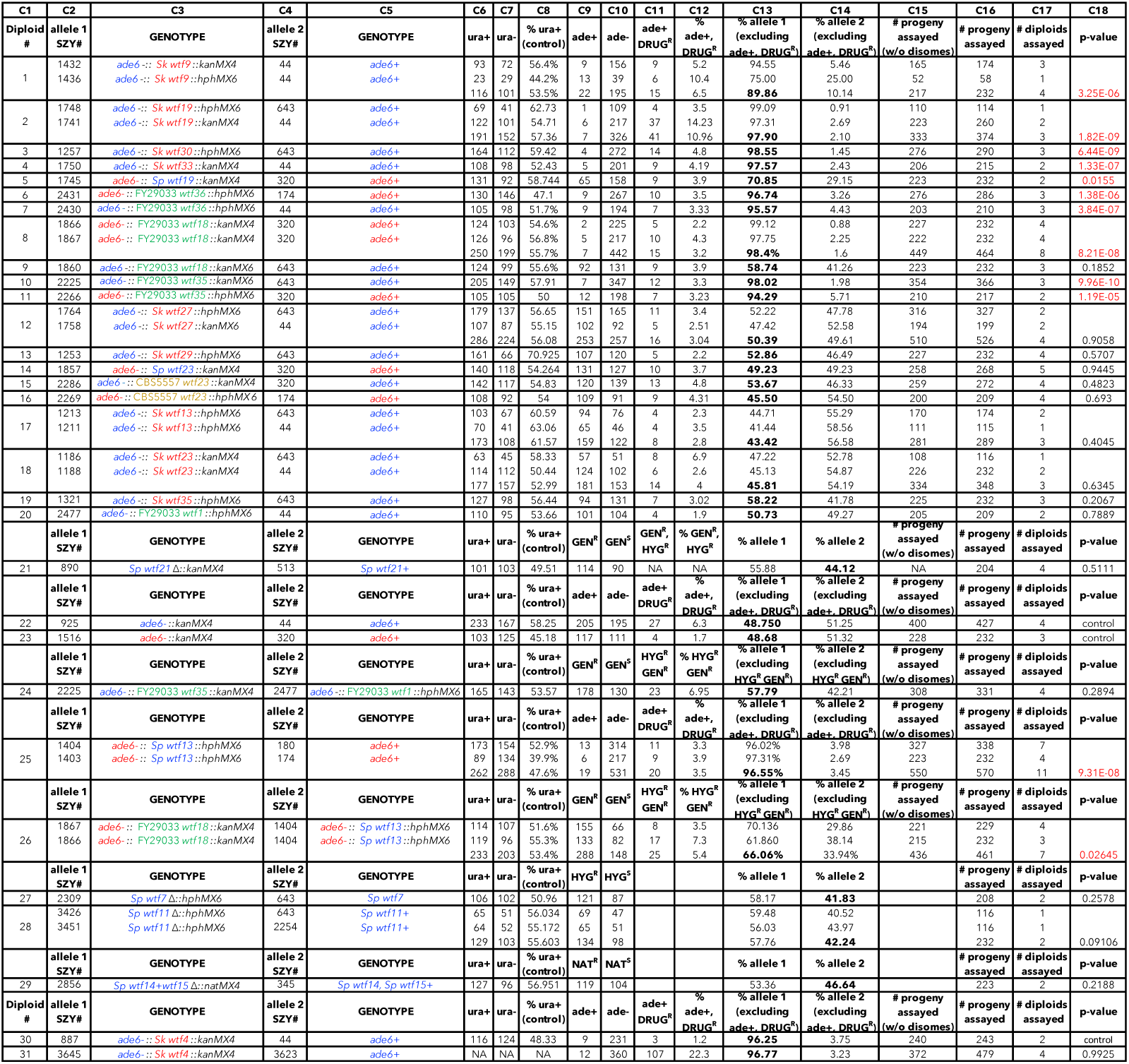
Raw data of allele transmission from Figure 2, 3, and Supplemental Figure 3. Each of the horizontal lines represents the relevant genotype and allele transmission from the indicated diploid into spores. The first column (C1) represents the diploid number, which matches the numbers in Figure 2, Figure 3, and Supplemental Figure 3. In columns C2-C5, the strain number (SZY) and relevant genotype of the haploid parent strains used to determine the allele transmission at the drive locus (*ade6* or *wtf* locus) are shown. *Sp, Sk,* CBS5557, and FY29033 alleles are labeled in blue, red, yellow, and green, respectively. Columns C6-C8 indicate which phenotypes were followed at the control locus (*ura4*) and the number of progeny that showed the indicated phenotypes. Columns C9 and C10 indicate the phenotypes that were followed at the drive loci (*ade6* or *wtf* locus) and the number of haploid progeny that exhibited the indicated phenotypes. Some of the progeny inherited both markers from the parent strains at the *ade6* locus. The number of the progeny that inherited both markers is presented in column C11 and the percentage of the progeny with this phenotype is shown in column C12. These progeny were excluded from the data presented in Figure 2, 3, and Supplemental Figure 3. Column C13 shows the fraction of the haploid progeny that inherited the genotype of allele 1. Column C14 shows the fraction of the haploid progeny that inherited the genotype of allele 2. Column C15 shows the total progeny assayed excluding the progeny that inherited both genetic markers at *ade6*. Column C16 shows the total progeny. Column C17 shows the total number of independent diploids assayed for each cross. The last column (C18) shows the p-value calculated by comparing diploids 1-4,7,9,10,12,13,15,17-20, and 24 to control diploid 22; diploids 5,6,8,11,14,16,25, and 26 to control diploid 23; diploids 21, 27, 28, and 29 to the control *ura4* locus; and diploid 31 to diploid 30. We previously published the allele transmission data for diploid 25 (18).

**Supplemental Table 2.** Raw data for fertility from Figure 3 and Supplemental Figure 3. Each column represents the diploid assayed, which matches the diploid number in Figure 3 and Supplemental Figure 3. The first row underneath the diploid number shows the SZY numbers of the haploid parent strains. We present all the viable spore yield values from independent assays. We normalized diploids 10, 20, and 24 to control diploid 22. We normalized diploids 8, 25, and 26 to diploid 23. We previously published the viable spore yield data for diploid 25 (18). We normalized the average viable spore yield value from the h^90^ mating of strain SZY3529 (*Sp wtf7*Δ, *wtf11*Δ, *wtf14+wtf15*Δ) to the average viable spore yield value of the wild-type control strain SZY2254. We calculated the p-value comparison using the Wilcoxon test.

| Figure 3 (VSY values) |  |  |  |  |  |  |  |  |  |  |  |
| --- | --- | --- | --- | --- | --- | --- | --- | --- | --- | --- | --- |
|  | Diploid 10 | Diploid 20 | Diploid 24 | Diploid 22 | Diploid 8 |  | Diploid 25 |  | Diploid 26 |  | Diploid 23 |
| haploid parents | 2225x643 | 2477x44 | 2225x2477 | 925x44 | 1866x320 | 1867x320 | 1404x180 | 1403x174 | 1866x1404 | 1867x1404 | 1516x320 |
|  | 1.393<br>1.718<br>1.675<br>1.121 | 1.700<br>1.424<br>2.000<br>2.419 | 3.553<br>2.380<br>3.455<br>2.064 | 2.217<br>2.106<br>2.733<br>2.200 | 0.857<br>0.932<br>1.582<br>0.776 | 1.157<br>1.406<br>1.121<br>1.070 | 1.229<br>1.050<br>3.000<br>2.667 | 2.378<br>1.796<br>1.625<br>1.850 | 1.411<br>2.870<br>1.368 | 2.175<br>2.153<br>2.774<br>1.741 | 3.256<br>2.982<br>1.930<br>4.154<br>3.778<br>3.216<br>3.189 |
| average | 1.477 | 1.886 | 2.863 | 2.314 | 1.113 |  | 1.949 |  | 2.070 |  | 3.215 |
| stdev | 0.277 | 0.426 | 0.752 | 0.284 | 0.273 |  | 0.683 |  | 0.604 |  | 0.696 |
| p-value | 0.02857 | 0.2 | 0.4857 | control | 0.0003108 |  | 0.00373 |  | 0.006993 |  | control |
| relative fertility | 63.81% | 81.48% | 123.70% | 100.00% | 34.61% |  | 60.63% |  | 64.40% |  | 100.00% |

**Supplemental Table 2. Raw data for fertility from Figure 3 and Supplemental Figure 3.**
| Supp Fig 3B (VSY values) |  |  |
| --- | --- | --- |
| haploid parent | 2254 | 3529 |
|  | 1.772<br>1.269<br>1.184<br>0.672<br>0.795<br>0.943 | 1.625<br>0.980<br>1.116<br>1.571<br>1.024 |
| average | 1.106 | 1.263 |
| stdev | 0.397 | 0.310 |
| p-value | control | 0.5368 |
| relative fertility | 100.00% | 114.24% |

**Supplemental Table 3.** Raw data of allele transmission from Supplemental Figure 4. Each of the horizontal lines represents the relevant genotype and allele transmission of the indicated diploid. The first column represents the diploid number, which matches the numbers in Supplemental Figure 4. In columns 2-5, the strain number (SZY) and relevant genotype of the haploid parent strains used to determine the allele transmission of chromosome 1 of *Sp* and *Sk*. Columns 6 and 7 indicate the number of progeny that exhibited the indicated phenotypes and were followed in chromosome 1 (*lys1* or *rec12*). Column 8 shows the total progeny assayed. Column 9 indicates the percent allele transmission of the *Sk* chromosome 1. Control diploid 32 (*Sk/Sk*) and diploid 34 (*Sp*/*Sk wtf1*Δ) are represented by one pair of haploid parents. Diploid 33 (*Sp/Sk* hybrid chromosome 1) shows the data from three different pairs of haploid parents. In two of the crosses, we followed chromosome 1 using the *lys1* markers. In the third cross, we followed chromosome 1 using the *rec12* markers. Column 10 shows the p-value (G-test) calculated by comparing diploid 33 and 34 to control diploid 32. Column 11 shows the total number of diploids assayed for each cross.

|  |  |  |  |  | chr1 allele transmission |  | # progeny assayed | % allele 2 | p value for % allele 2 | # diploids assayed |
| --- | --- | --- | --- | --- | --- | --- | --- | --- | --- | --- |
| Diploid # | allele 1 SZY# | GENOTYPE | allele 2 SZY# | GENOTYPE | lys+ | lys- |  |  |  |  |
| **32** | 293 | *lys1+* | 298 | *lys1 Δ::kanMX4* | 211 | 232 | 443 | **52.37** | control | 2 |
| **33** | 3517 | *lys1+* | 196 | *lys1 Δ::kanMX4* | 181 | 283 | 464 | 60.99 |  | 4 |
|  | 3518 | *lys1+* | 196 | *lys1 Δ::kanMX4* | 126 | 222 | 348 | 63.79 |  | 4 |
|  | allele 1 SZY# | GENOTYPE | allele 2 SZY# | GENOTYPE | ura- | ura+ |  |  |  |  |
|  | 147 | *rec12-117* | 298 | *rec12 Δ::ura4+* | 257 | 511 | 768 | 66.54 |  | 3 |
|  | **Total Diploid 33** |  |  |  | 564 | 1016 | 1580 | **64.30** | **0.02243** | 11 |
| **34** | 147 | *rec12-117* | 3829 | *rec12 Δ::ura4+, wtf1 Δ* | 354 | 690 | 1044 | **66.09** | **0.01352** | 11 |

**Supplemental Table 4.**
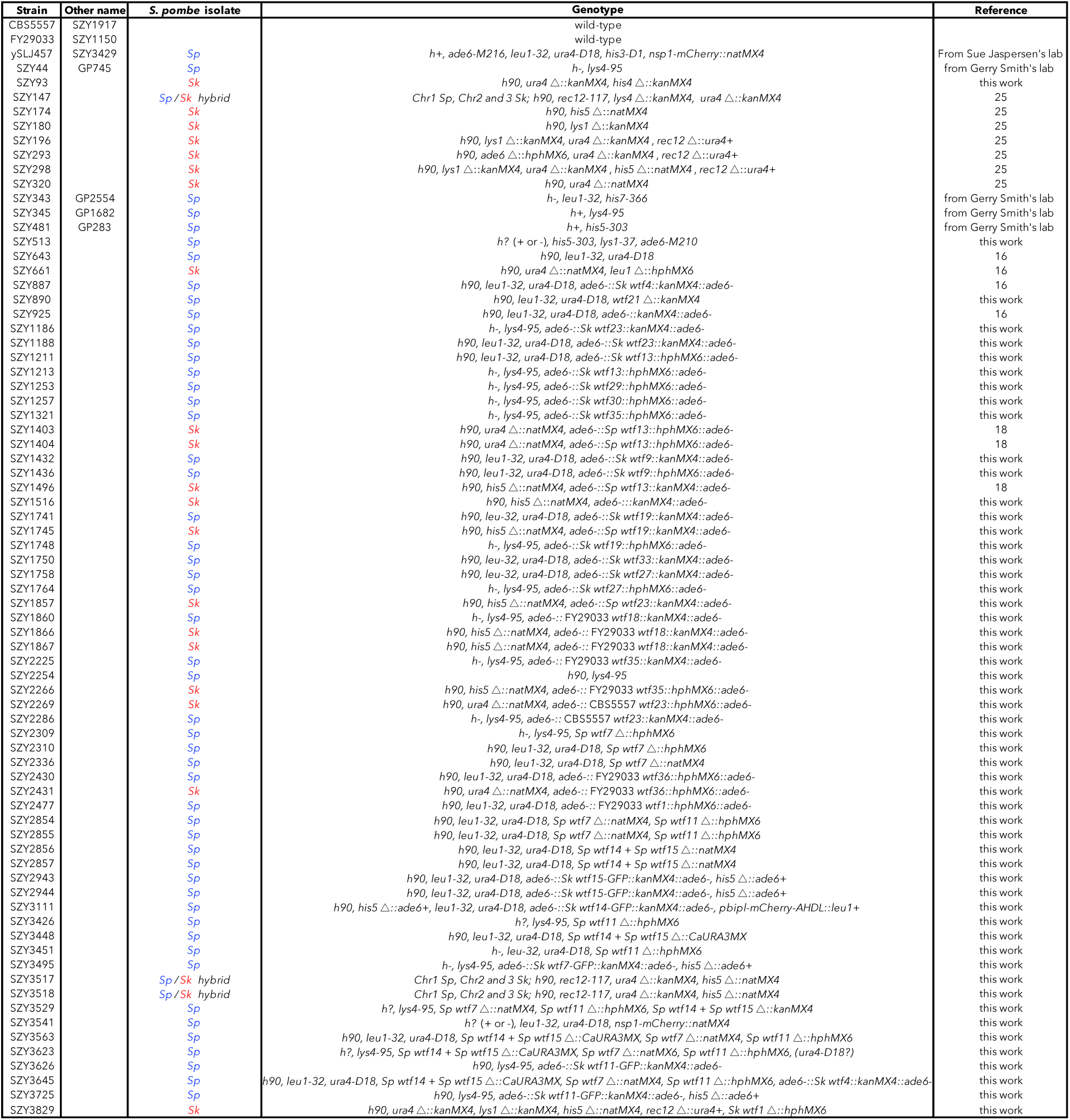
Yeast strains used in this study. The strain SZY643 contains the *wtf18-2* allele (18).

**Supplemental Table 5.** Table summary of plasmid constructions. Column 1 lists the *wtf* gene cloned into each vector. Column 2 denotes the isolate origin of each *wtf* gene in column 1. The DNA templates and oligos used in the PCR reactions to amplify the *wtf* alleles are shown in columns 3 and 4, respectively. We digested each of the amplified fragments with the enzymes reported in column 5 and then integrated into the target site listed in column 6. The number of each of the plasmids that we generated is reported in column 7. The description of each plasmid can be found in Supplemental Table 6.

| *wtf* allele | S. pombe isolate | DNA template (gDNA) | PCR oligos used | PCR fragment digested with restriction enzyme | target site | Resulting plasmid number |
| --- | --- | --- | --- | --- | --- | --- |
| *wtf19* | *Sp* | SZY643 | 1129 + 1130 | SacI | SacI site of pSZB188 | pSZB507 |
| *wtf23* | *Sp* | SZY44 | 890 + 891 | SacI | SacI site of pSZB188 | pSZB372 |
| *wtf13* | *Sk* | SZY180 | 2260+2261 | SpeI | SpeI site of pSZB386 | pSZB399 |
| *wtf14* | *Sk* | SZY180 | 881+882 | SacI | SacI site of pSZB188 | pSZB378 |
| *wtf19* | *Sk* | SZY661 | 1129+1133 | SacI | SacI site of pSZB188 | pSZB511 |
| *wtf19* | *Sk* | SZY661 | 1129+1133 | SacI | SacI site of pSZB386 | pSZB512 |
| *wtf23* | *Sk* | SZY180 | 890 + 891 | SacI | SacI site of pSZB188 | pSZB379 |
| *wtf27* | *Sk* | SZY661 | 1134+1135 | SpeI | SpeI site of pSZB188 | pSZB516 |
| *wtf27* | *Sk* | SZY661 | 1134+1135 | SpeI | SpeI site of pSZB386 | pSZB519 |
| *wtf30* | *Sk* | SZY180 | 977+978 | SpeI | SpeI site of pSZB386 | pSZB410 |
| *wtf33* | *Sk* | SZY661 | 1131+1132 | SacI | SacI site of pSZB188 | pSZB514 |
| *wtf35* | FY29033 | FY29033 | 1036+1349 | SacI | SacI site of pSZB188 | pSZB788 |
| *wtf35* | FY29033 | FY29033 | 1036+1349 | SacI | SacI site of pSZB387 | pSZB800 |
| *wtf36* | FY29033 | FY29033 | 1351+1593 | SacI | SacI site of pSZB387 | pSZB852, pSZB853 |
| *wtf23* | CBS5557 | CBS5557 | 890+891 | SacI | SacI site of pSZB188 | pSZB810 |
| *wtf23* | CBS5557 | CBS5557 | 890+891 | SacI | SacI site of pSZB387 | pSZB812 |

**Supplemental Table 6.** Plasmids used in this study.

| Plasmids | short description | reference |
| --- | --- | --- |
| pFA6 | contains *kanMX4* | 56 |
| pAG25 | contains *natMX4* | 57 |
| pAG32 | contains *hphMX6* | 57 |
| pKT127 | contains yEGFP | 61 |
| pFA6-mTurq2-URA3MX | contains mTurq2 and CaURA3MX | 59 |
| pMZ222 | pART1 with Cas9 (*adh1* promoter) | 60 |
| pMZ283 | pUR19 with *rrk1*-gRNA (CspCI targeting sequence placeholder) | 60 |
| pSZB188 | derivative of pFA6 that integrates at *ade6*, yielding *ade6*- | 16 |
| pSZB189 | pSZB188 with *Sk wtf4* cloned into SacI site | 16 |
| pSZB197 | pMZ283 with gRNA targeting *wtf21* cloned into the CspCI site | this work |
| pSZB372 | pSZB188 with *Sp wtf23* cloned into SacI site | this work |
| pSZB378 | pSZB188 with *Sk wtf14* cloned into SacI site | this work |
| pSZB379 | pSZB188 with *Sk wtf23* cloned into SacI site | this work |
| pSZB386 | derivative of pAG32 that integrates at *ade6*, yielding *ade6*- | 18 |
| pSZB387 | derivative of pAG32 that integrates at *ade6*, yielding *ade6*- | 18 |
| pSZB399 | pSZB386 with *Sk wtf13* cloned into SpeI site | this work |
| pSZB410 | pSZB386 with *Sk wtf30* cloned into SpeI site | this work |
| pSZB507 | pSZB188 with *Sp wtf19* cloned into SacI site | this work |
| pSZB511 | pSZB188 with *Sk wtf19* cloned into SacI site | this work |
| pSZB512 | pSZB386 with *Sk wtf19* cloned into SacI site | this work |
| pSZB514 | pSZB188 with *Sk wtf33* cloned into SacI site | this work |
| pSZB516 | pSZB188 with *Sk wtf27* cloned into SpeI site | this work |
| pSZB519 | pSZB386 with *Sk wtf27* cloned into SpeI site | this work |
| pSZB691 | pSZB188 with *Sk wtf7-GFP* cloned into SacI site | this work |
| pSZB696 | pSZB188 with *Sk wtf14-GFP* cloned into SacI site | this work |
| pSZB698 | pSZB188 with *Sk wtf15-GFP* cloned into SacI site | this work |
| pSZB788 | pSZB188 with FY29033 *wtf35* cloned into SacI site | this work |
| pSZB800 | pSZB387 with FY29033 *wtf35* cloned into SacI site | this work |
| pSZB810 | pSZB188 with CBS5557 *wtf23* cloned into SacI site | this work |
| pSZB812 | pSZB387 with CBS5557 *wtf23* cloned into SacI site | this work |
| pSZB852 | pSZB387 with FY29033 *wtf36* cloned into SacI site | this work |
| pSZB853 | pSZB387 with FY29033 *wtf36* cloned into SacI site | this work |
| pSZB879 | pSZB387 with FY29033 *wtf1* cloned into SacI site | this work |
| pSZB1087 | pSZB188 with *Sk wtf11-GFP* cloned into SacI site | this work |

**Supplemental Table 7.**
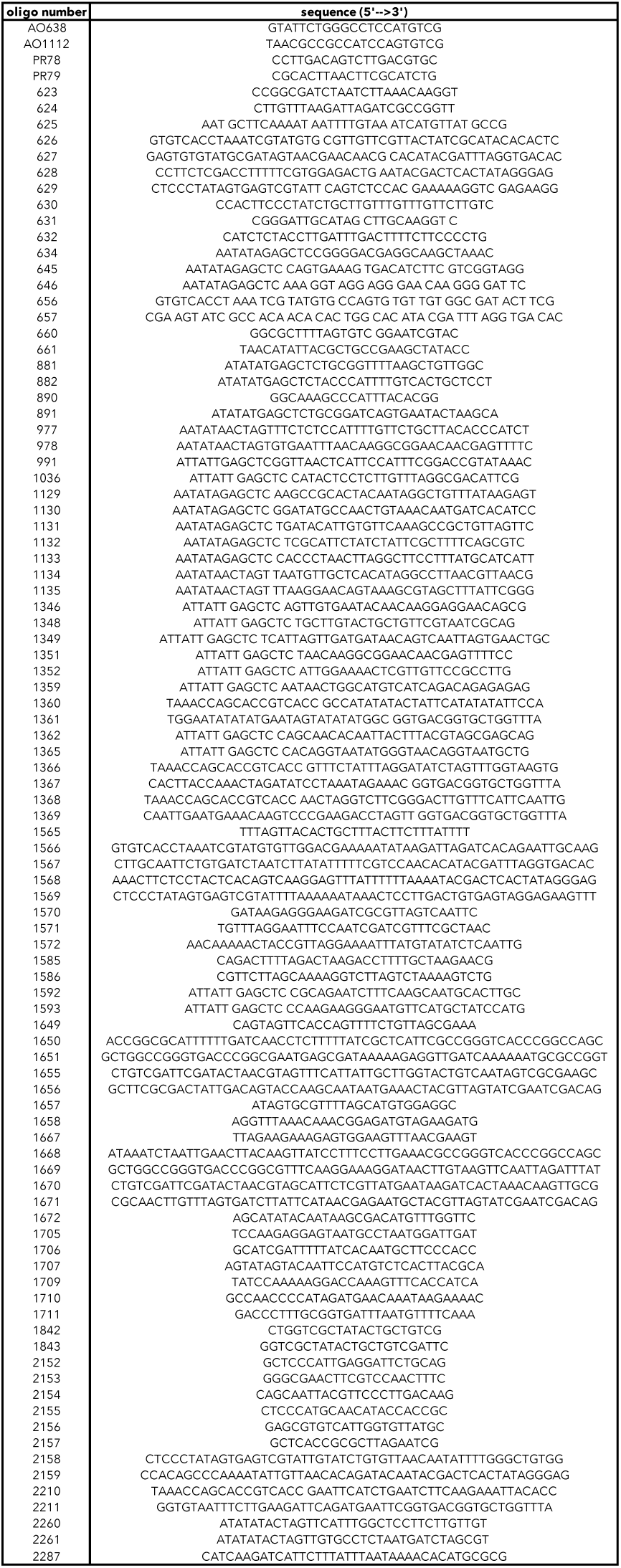
Oligos used in this study.

